# Single nucleus analysis of Arabidopsis seeds reveals new cell types and imprinting dynamics

**DOI:** 10.1101/2020.08.25.267476

**Authors:** Colette L. Picard, Rebecca A. Povilus, Ben P. Williams, Mary Gehring

## Abstract

Seeds are the basis of agriculture, yet their full transcriptional complexity has remained unknown. Here, we employ single-nucleus RNA-sequencing to characterize developing *Arabidopsis thaliana* seeds, with a focus on endosperm. Endosperm, the site of gene imprinting in plants, mediates the relationship between the maternal parent and embryo. We identify new cell types in the chalazal endosperm region, which interfaces with maternal tissue for nutrient unloading. We further demonstrate that the extent of parental bias of maternally expressed imprinted genes varies with cell cycle phase, and that imprinting of paternally expressed imprinted genes is strongest in chalazal endosperm. These data indicate imprinting in endosperm is heterogeneous and suggest that parental conflict, which is proposed to drive the evolution of imprinting, is fiercest at the boundary between filial and maternal tissues.

---

As nutrient-filled units of propagation, seeds are a key life-cycle stage for many plants and are the primary source of calories consumed by humans (*1*). Flowering plant seeds are uniquely complex structures, being comprised of a diploid maternally-derived seed coat that surrounds two products of distinct fertilization events – the embryo and endosperm. The embryo represents the next diploid generation of the plant life cycle. The endosperm is an often triploid tissue (due to the presence of an additional maternal genome complement), and is an altruistic mediator of the relationship between its sibling embryo and their resource-supplying mother. Endosperm is a key evolutionary innovation of flowering plants and has been identified as the site of genomic imprinting, an epigenetic gene regulatory process that results in differential expression of maternally and paternally inherited alleles (*1,2*). Although an ephemeral tissue, endosperm undergoes a unique developmental program that includes differentiation into three morphologically and spatially-defined domains: the micropylar domain surrounds the embryo, the chalazal domain occupies the opposite end of the seed, and the peripheral domain lies in between (*3–6*). Gene expression patterns in the three endosperm domains have been assessed by microarray analysis (*7*), but it is unknown whether cell-type heterogeneity exists within domains. Despite its evolutionary and agronomic importance, endosperm biology remains relatively little understood. A complete record of all transcriptionally unique cell or nuclei types within the endosperm has been unobtainable due to the compact, interconnected, and complex nature of seeds.

To build a comprehensive map of transcriptional complexity and to examine imprinting dynamics during early endosperm development in Arabidopsis, we performed single nucleus RNA-seq (snRNA-seq). We isolated nuclei instead of cells because the endosperm is syncytial during its early development, before undergoing progressive cellularization. Most profiled nuclei were from F_1_ seed from reciprocal crosses between the wild-type strains Col and Cvi collected at 4 days after pollination (DAP) (Fig. 1A). We obtained high-quality transcriptomes for 1,437 nuclei using fluorescence-activated sorting of DAPI-stained seed nuclei to enrich for 3C/6C endosperm nuclei, combined with a modified smart-seq2 protocol (*8*) for library preparation (Fig. 1A, Figs. S1-S3). We used the SC3 program (*9*) to cluster all snRNA-seq data, obtaining 27 clusters ranging in size from 8 to 172 nuclei (Fig. S4). Based on initial clustering and the fraction of maternal allele expression per nucleus, we identified 966 endosperm nuclei, 464 seed coat nuclei, and 7 embryo nuclei (Fig. S1, Fig. S4, Supplementary Materials).

**Fig. 1.**
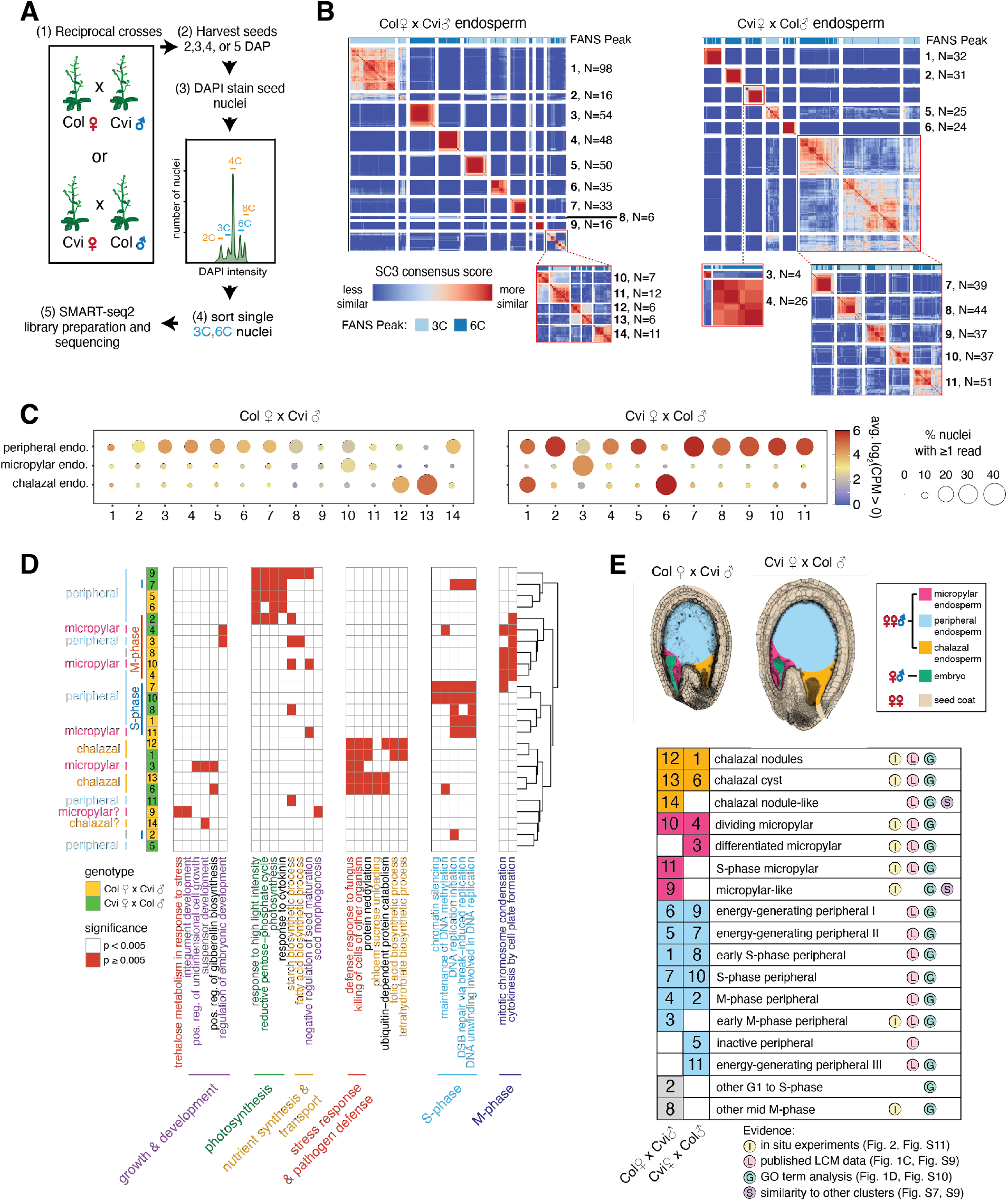
Distinct nuclei types in Arabidopsis endosperm. (A) Overview of experimental approach. (B) SC3 clustering of Col x Cvi and Cvi x Col 4 DAP endosperm nuclei. Insets: re-clustering to further resolve distinct groups. (C) Average expression of marker genes for peripheral, micropylar, and chalazal endosperm regions, based on (*7,27*). (D) Heatmap of a subset of significantly enriched gene ontology terms among genes upregulated in each cluster. (E) Seed images at 4 DAP, with seed regions false-colored, and identification of the nuclei states corresponding to each cluster.

To test whether our clustering strategy reliably identified distinct cell or nuclei types, we took advantage of the 343 seed coat nuclei collected at 4 DAP, which were captured due to the proximity of the 3C and 4C peaks (Fig. S1). The seed coat is comprised of six distinct cell layers or regions (*7,10*). Clustering yielded 6 clusters for Col-derived seed coat (from Col x Cvi crosses) and 8 clusters for Cvi-derived seed coat (from Cvi x Col crosses) (Fig. S5). To assign putative identities to the computationally-defined clusters, we evaluated the expression of known seed-coat cell layer markers and performed GO term enrichment analysis on differentially expressed genes (Fig. S5-S9). Our clustering and characterization recovered all known seed coat cell types and provides the first whole-genome expression data for distinct layers and regions of the seed coat (Fig. S5, Supplementary Material).

We next applied our analysis method to the 966 endosperm nuclei. A single Arabidopsis seed has ~350 endosperm nuclei at the stage assayed (*11*), so this dataset should represent a near complete sampling. We identified 14 distinct clusters of nuclei in Col x Cvi F_1_ endosperm (CxV E1-E14) and 11 clusters in Cvi x Col (VxC E1-E11) (Fig. 1B), suggesting there is cell identity heterogeneity within the three known endosperm domains. We determined the identity of endosperm clusters by evaluating the expression of known marker genes for micropylar, peripheral, chalazal endosperm, differential gene expression and GO term enrichment analysis, *in situ* hybridization for cluster-specific transcripts, and cell cycle trajectory analysis. We identified several endosperm clusters corresponding to micropylar and peripheral endosperm nuclei, some related to cell cycle phase and others to functional differences (Fig. 1D,E, Fig. 2, Fig. S7, Fig. S9-S14, Supplementary Material).

**Fig. 2.**
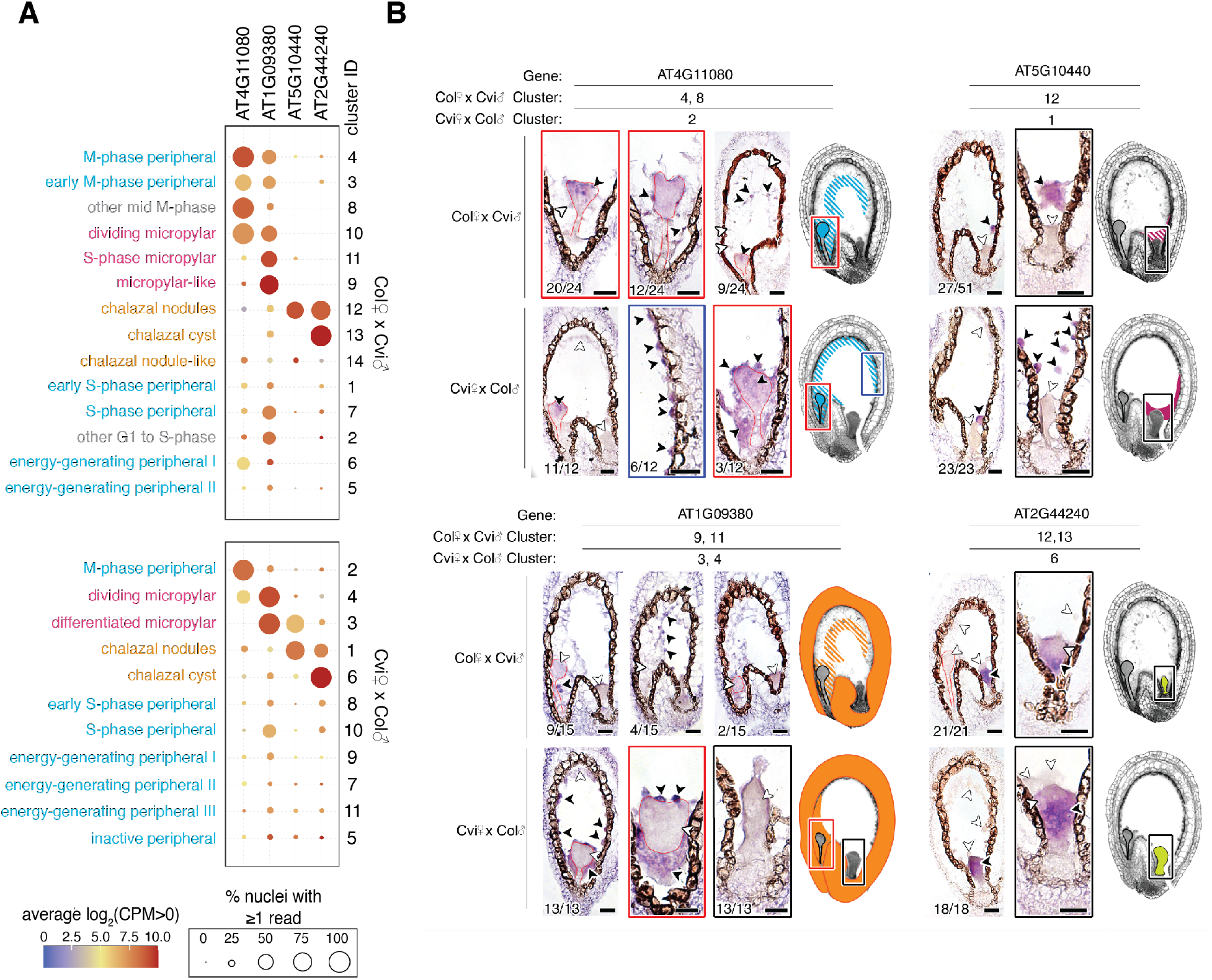
Identification of clusters by *in situ* hybridization analysis. (A) Average expression of four cluster-specific genes selected for *in situ* hybridization. (B) RNA *in situ* hybridization in 4 DAP seeds. False-colored images summarize gene expression patterns for each locus and cross direction. Solid colors, consistent expression; striped pattern, variable expression. Black arrowheads, transcript detected; white arrowheads, no transcript detected. Embryos outlined in red. Number of seeds with the pictured expression pattern, as well as total number of seeds observed, indicated in bottom left of each image. Images without seed numbers represent higher magnification images of an already presented expression pattern. Sense probe images and hybridization for additional transcripts in Fig. S11. Scale bars, 25 μm.

Gene expression analysis and the overlap of known endosperm domain markers suggested that at least two distinct clusters in each genotype corresponded to chalazal endosperm, which is thought to be a primary site of nutrient transfer between the mother and offspring (*12*) (Fig. 1C,D, Fig. S7, Fig. S9). This was unexpected because no transcriptional or functional distinctions within chalazal endosperm have been described. However, anatomically, the chalazal endosperm consists of two regions: the chalazal nodules are large, possibly multinucleate bodies lining the chalazal region (*13*), while the chalazal cyst is a cytoplasmically-dense, multinucleate region that forms at the interface between the endosperm and adjacent maternal tissue (*14*). RNA *in situ* hybridization analysis showed that two transcripts most highly expressed in CxV E12 and VxC E1, AT1G44090 and AT5G10440, were localized specifically to the chalazal nodules (Fig. 2, Fig. S11). In contrast, AT2G44240 and AT4G13380, which are primarily expressed in CxV E13 and VxC E6, were only detected in the chalazal cyst by *in situ* hybridization (Fig. 2, Fig. S11). We concluded that CxV E12 and VxC E1 correspond to the chalazal nodules, while CxV E13 and VxC E6 correspond to the cyst (Fig. 1E, Fig. 2B). Remarkably, despite the lack of cell membranes and walls in chalazal endosperm, physically adjacent nodule and cyst cytoplasmic domains did not share the expression of cluster-specific genes (Fig. 2B).

Cell cycle phase further distinguished the chalazal cyst and nodules. Chalazal nuclei as a whole were predominantly in G1, G1/S, S and G2, but rarely in M phase, suggesting they undergo endoreduplication (Fig. S12,13). This is consistent with observations that chalazal endosperm nuclei are larger than other endosperm nuclei and likely polyploid (*6,14*), and with our finding that chalazal nuclei were preferentially sorted from the 6C peak (Fig. 1B). More than half of nodule nuclei were in G1/S or S-phase, while most cyst nuclei were in G1 or G2 (Fig. S12). No M phase nuclei were detected in the chalazal cyst. Thus, the cyst consists primarily of nuclei that are non-dividing or that spend little time in S-phase.

All chalazal clusters showed high expression of genes related to pathogen defense and cell killing, as well as protein neddylation (Fig. 1D, Fig. S10). Additionally, genes highly expressed in chalazal nodules were involved in tetrahydrofolate and folic acid biosynthesis, a key step in one-carbon metabolism and a major target process for crop biofortification (*15*). By contrast, the cyst was enriched for ubiquitin-dependent protein catabolism and phloem sucrose unloading (Fig. 1D). The chalazal cyst is adjacent to the termination of maternal phloem tissue in the chalazal seed coat region, and the enrichment of genes related to phloem sucrose unloading is consistent with a nutrient transfer function for the cyst. Taken together, these experiments indicate that chalazal endosperm consists of two transcriptionally, spatially, developmentally, and functionally distinct nuclei types: nodules and cysts.

We next took advantage of the allele-specific nature of our data to examine imprinted expression across the endosperm nuclei clusters we defined. Investigation of parental bias in endosperm allele-specific bulk mRNA-seq datasets (*16–21*) demonstrate that while imprinted genes are, by definition, significantly biased toward expression from either the maternal or paternal allele, few are expressed monoallelically. Partial imprinting could result from incomplete silencing of the non-expressed allele throughout the endosperm, or from heterogeneous imprinting among individual cells or cell types. Understanding whether endosperm imprinting is heterogenous is important for understanding both the cellular and physiological function of imprinting and its underlying epigenetic basis.

We developed a novel analysis framework for evaluating genomic imprinting from snRNA-seq data and one that is suitable for situations where maternal (m) and paternal (p) allelic dosage is not 1:1 (endosperm is 2m:1p) (Fig. S15-S17, Supplementary Material). We detected significant maternal bias for 357 genes and paternal bias for 110 genes (Fig. S18-20, Data S4). MEGs and PEGs were defined as strong, medium, or weak based on the extent of parental bias (Fig. S18,19).

To determine whether imprinted genes were preferentially expressed in specific nuclei types within endosperm, we examined total and allelic expression patterns for these genes across endosperm clusters. MEG expression was not enriched in any specific endosperm nuclei type, with a few exceptions for individual genes (Fig. S21). By contrast, nearly half of the PEGs we identified had strongly enriched expression in the chalazal endosperm clusters (Fig. 3A,B, Fig. S21). A subset of these was specifically enriched in the chalazal nodules, while another subset was enriched in the cyst. The increased expression of PEGs in chalazal endosperm was caused by increased paternal allele expression, while maternal allele expression remained low and largely unchanging across all endosperm clusters (Fig. 3B,C; Fig. S22). This effect was not observed for non-imprinted genes with enriched chalazal expression (Fig. S23). Thus, the paternal allele of many PEGs becomes specifically upregulated in the chalazal endosperm region. Taken together, these results demonstrate that imprinting is heterogenous among endosperm cell types.

**Fig. 3.**
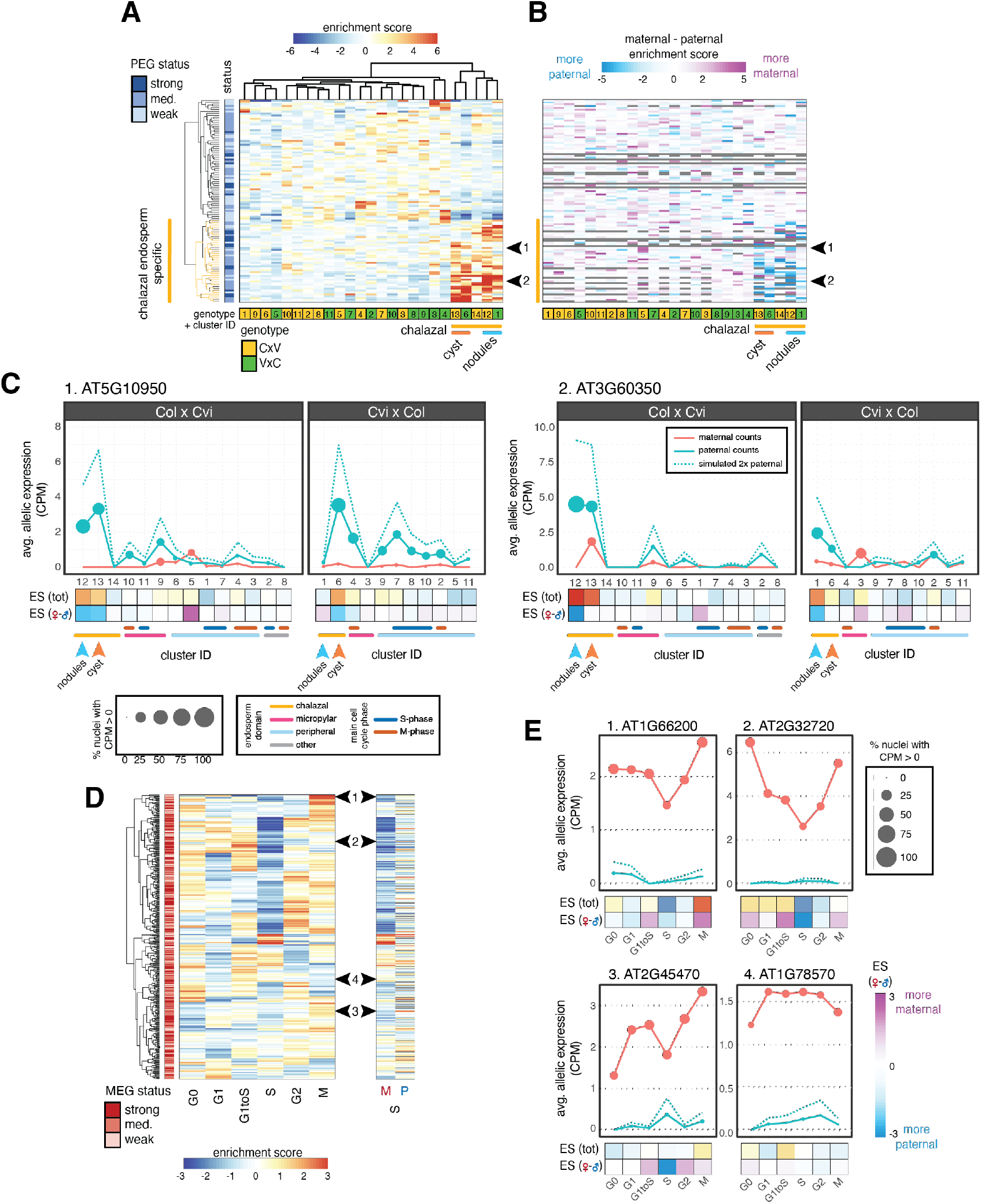
Imprinting heterogeneity in endosperm. (A) A large fraction of PEGs are specifically expressed in chalazal endosperm. Heatmap of total expression enrichment scores (ES) for all PEGs. (B) Heatmap of ES (maternal) - ES (paternal), the difference between the allele-specific maternal and paternal expression ES. (C) Average allelic expression of nuclei in Col x Cvi and Cvi x Col endosperm clusters for two example PEGs, indicated by black arrows in (A) and (B). Dotted blue line represents simulated expression from two paternal genomes. (D) Heatmap of MEG total expression ES (left), maternal (M) and paternal (P) allele-specific ES for S-phase (right). Row order same for all heatmaps. (E) Average allelic expression for three MEGs that show reduced maternal allele expression in S-phase along with one MEG that does not, indicated by black arrows in (D). CPM, counts per million.

Imprinted gene expression is regulated epigenetically, with DNA methylation and the PRC2 histone mark H3K27me3 playing important roles in regulating differential allelic expression (*22–24*). We therefore asked whether known epigenetic regulators were differentially expressed in chalazal nuclei. Genes downregulated in the chalazal nodule clusters were enriched for the GO term ‘regulation of genomic imprinting’ because of reduced expression of the PRC2 gene *FIE*, the DNA maintenance methyltransferase *MET1*, and the 5-methylcytosine DNA glycosylase gene *DME* (Fig. S10, Fig. S24). However, other epigenetic regulators were upregulated. The PRC2 histone methyltransferase *MEA* and the histone H3K4 demethylase *JMJ15*, which are both MEGs, were specifically expressed in the chalazal nodules (Fig. S24). The *Su(var)3-9* homologues *SUVH7* (a PEG) and *SUVH8* were specific to the chalazal cyst, as was the 5-methylcytosine DNA glycosylase *DML3* (Fig. S24). Other epigenetic regulators were highly expressed in both nodule and cysts but not in other nuclei types (Fig. S24). These data suggest that targeted up- and down-regulation of specific epigenetic regulators may help create a distinct epigenetic state in the chalazal region compared to other endosperm regions, resulting in upregulation of the paternal allele of PEGs in these nuclei. Some of these epigenetic regulators are known to regulate imprinted expression (*23*), suggesting these factors may be mediating an active parental conflict within the chalazal nodules, perhaps by opposing or promoting elevated expression of PEGs. Alternatively, a chalazal endosperm-specific transcription factor could interact with differential maternal and paternal allele epigenetic states to specifically promote expression of the paternal allele of PEGs.

Our dataset also allowed us to examine MEG and PEG expression patterns as a function of the cell cycle, which has not been systematically assayed in either mammals or flowering plants. Expression of nearly half of the MEGs identified in our analysis decreased during S-phase (Fig. 3D, Fig. S25). This pattern was not observed for PEGs or for a set of 500 randomly selected, non-imprinted genes (Fig. S22). The lower S-phase expression of MEGs was associated with decreased maternal bias of MEGs, caused by reduced expression of the maternal allele (Fig. 3D,E; Fig. S25). During S-phase, chromatin states are disrupted and reassembled as DNA is replicated. These data suggest that MEG expression may be particularly sensitive to disruptions in epigenetic state that transiently occur during DNA replication.

We have shown that the endosperm of *A. thaliana* contains a previously undescribed diversity of transcriptionally and functionally distinct cell/nuclei types. One important conclusion from this work is that for a subset of imprinted genes, imprinting is dynamic across the cell cycle and/or heterogenous between cell/nuclei types. In particular, many PEGs are most strongly paternally biased in the chalazal endosperm region. This is especially noteworthy in light of the theory that imprinting evolved in flowering plants and mammals as an outcome of conflicts between parental genomes in asymmetrically related offspring over maternal resource transfer (*25,26*). The high expression of paternal alleles of PEGs in chalazal endosperm suggests that this conflict is strongest at the interface between maternal and filial tissues in developing seeds. Our study further suggests that fully understanding the regulatory mechanisms underlying imprinting will require cell/nuclei-type specific approaches.

## Acknowledgements

We thank the MIT BioMicro Center and the Whitehead Institute Genome Technology Core and Flow Cytometry Core Facility for research assistance, and Francine Lafontaine for valuable input on statistical methods.

## Author contributions

C.L.P. and M.G. conceived the project; C.L.P. and R.A.P. conducted experiments; C.L.P., R.A.P., and B.W. analyzed data with input from M.G., C.L.P. and M.G. wrote the manuscript with edits from R.A.P and B.P.W.

