## Supplementary Material for "Single nucleus analysis of Arabidopsis seeds reveals new cell types and imprinting dynamics"

Materials and Methods

Supplementary Text

Figs. S1-S25

**Materials and Methods**

Plant material and crossing

All Col-0, L*er* and Cvi-0 parent plants were grown in a growth chamber under 16h light at 50% relative humidity (16h at 22°C and 120µm light, 8h at 20°C and 0µm light). Plants were emasculated in the afternoon or evening, and pollinated in the morning two days later. FANS was performed in the morning to maximize consistency in seed stage across experiments. However, different crosses developed at different rates: the endosperm of the average Col x Cvi (CxV) $F_1$ seed (mother in cross is indicated first) had already begun to cellularize at 4 DAP, while Cvi x Col (VxC) $F_1$ seeds were generally still in the proliferative phase at 4 DAP (Fig. 1E). Embryo developmental stage at 4 DAP was also more variable in CxV crosses, whereas most 4 DAP VxC seeds were at the heart stage (Fig. S1). VxC seeds are larger than CxV seeds (Fig. 1E).

Confocal microscopy

Controlled floral pollinations were performed for each cross depicted in Fig. 1E. Siliques were collected 4 DAP, dissected to open up the carpel wall, and fixed in FAA overnight at 4°C. Samples were dehydrated with an ethanol series to 100% ethanol, and cleared by gradual infiltration with immersion oil (Immersol 518F, Zeiss). Seeds were removed from cleared siliques and mounted on slides. Whole seeds were imaged on a Zeiss LSM700 Confocal Microscope, with settings optimized to detect auto-fluorescence (Excitation: 405nm at 0.1% power, 488nm 11% power, 555nm at 28% power. Detection: from 415 to 735 nm.).

RNA *in situ* hybridizations

Controlled floral pollinations were performed for each cross; more than 10 cross pollinations were performed per cross type. Siliques were harvested 4 DAP and fixed in FAA overnight at 4°C. Following dehydration and clearing (HistoClear, National Diagnostics), samples were embedded in Paraplast Plus (McCormick Scientific) with vacuum infiltration, and sectioned at 8 µM (Leica RM 2065 rotary microtome). Ribbons were mounted with DEPC water on ProbeOn Plus slides (Fisher) at 42°C and dried overnight at 37°C. For probes against marker genes, primers and fragments sizes are listed below, and the previously published 602 bp probe for *PDF1* was used as a positive control (*28*).

| Name | Gene ID | Length (bp) | Forward Primer (5'-3') | Reverse Primer( 5'-3') |
|---|---|---|---|---|
| CYCD4;2 | AT5G10440 | 437 | ACCAAAGCAAATTCTACAATAGAG | GTCCAAATTGAAGTTCCTCACAG |
| GA20OX5 | AT1G44090 | 530 | TAGGCAACCATCGTCAAGAGATC | CCTGTGGTAACAACTCCTGTAG |
| MEE56 | AT4G13380 | 472 | AGGCAGAAATCTTGACGATGAATG | GCCATTTCTTTCTCATCATCACTAC |
| UMAMIT25 | AT1G09380 | 585 | AGGCATCAGCACAAGCCAAAG | ATACGAACTAGAGACCAATAACCAG |
| HMG1,2 | AT4G11080 | 634 | GACGGAGCTGAAGAAACTGC | CGCCATCTGATCATAAGGAGCC |
| NEP-interacting | AT2G44240 | 622 | GGTTTTGCGGTAGCTCTGATG | GATAAACCTGCCAACCAGCTTA |

Probes were amplified from endosperm cDNA and cloned into TOPO pCR II or TOPO pCR 4 vectors (ThermoFisher). Plasmids containing sense and antisense oriented fragments were identified and linear templates were amplified using M13 forward and reverse primers for probe synthesis. Antisense and sense RNA probes were synthesized in vitro with digoxigenin-UTPs using T7 or SP6 polymerase (DIG RNA labeling kit, Roche/Sigma-Aldrich). Probes were subsequently hydrolyzed to approximately 300 bp and dot blots were performed to estimate probe concentration. Pre-hybridization steps were preformed according to (*29*), except Pronase digestion occurred for 15 minutes at 37°C. Hybridization and post-hybridizations were performed according to (*30*), with minor modifications. For higher confidence in directly comparing expression patterns, slides corresponding to the cross and its reciprocal were processed face to face in the same pairs for hybridization, antibody, and detection steps. Negative controls consisted of hybridizing sense probes to tissue from each cross direction (Fig. S11). Hybridization was performed overnight at 55°C, slides were then washed twice in 0.2X SSC for 60 mins each at 55°C, then twice in NTE for 5 min at 37°C and RNaseA treated for 20 min at 37°C, followed by two more 5 min NTE washes. Slides were incubated at room temperature for 1 hour with Anti-DIG antibody (Roche/Sigma Aldrich) diluted 1:1250 in buffer A (*30*). Slides were then washed four times for 20 min each at room temperature with buffer A and once for 5 min with detection buffer (*30*). Colorimetric detections were performed using NBT/BCIP Ready-To-Use Tablets (Roche/Sigma-Aldrich) dissolved in water or BM-Purple (Sigma-Alrdich) with Levamisole (Vector Laboratories). Slides were allowed to develop 16–46 hours before stopping color precipitation by washing briefly with 50% and then 100% ethanol (NBT/BCIP) or 50% and then 100% methanol (BM Purple). Slides were mounted in Permount (Electron Microscopy Sciences) and imaged using a Zeiss Axio Imager M2. Color and brightness/contrast adjustments and smart sharpen were applied to whole images, with particular attention to having even white-balance across different images (Adobe Photoshop).

<u>Seed nuclei FANS</u>
Since the endosperm is a syncytium at most of the timepoints used in this study, and nuclei transcriptomes are well-correlated with whole-cell transcriptomes (*31*), we isolated nuclei instead of cells. For fluorescent activated nuclei sorting (FANS), seeds were manually removed from siliques (~ 2 siliques per sample) into 50 μL Partec nuclei extraction buffer. Samples were disrupted using a blue pestle in a microfuge tube before adding 400 μL Partec nuclei staining buffer and mixing by pipetting. Samples were filtered twice through a 30μm nylon mesh (Partec CellTrics #04-004-2326, Sysmex). For samples sorted on 9/12/2018, 9/13/2018, 9/20/2018 and 9/26/2018, two additional wash steps were performed to potentially remove cell lysate from the sample. For each wash, nuclei were spun down 5 min at 1000 g in a centrifuge pre-cooled to 4°C. Supernatant was then removed and nuclei were gently resuspended in 1 mL of a 1:8 mix of Partec nuclei extraction buffer and Partec nuclei staining buffer. Individual nuclei were sorted into wells a 96-well PCR plate. A total of 22 full or partial plates (batches) of samples were prepared. Each plate included at least one negative control (no nucleus sorted into well) and one positive control (50 nuclei sorted into a single well); however, these controls were not sequenced in the first batch. Some plates also included wells with 2 nuclei sorted into each as controls for the precision of single-nuclei sorting. For most sorting experiments, a small number of seeds were separately cleared with a chloral hydrate buffer and imaged in order to determine seed developmental stage (Fig. S1). Nuclei were sorted from both the putative 3C and 6C ploidy peaks based on DAPI fluorescence to enrich for endosperm nuclei (see Fig. S1).

<u>snRNA-seq library preparation and sequencing</u>
FANS samples were prepared as outlined above in the morning either 2, 3, 4 or 5 days after pollination (DAP). Libraries were prepared according the Smart-seq v2 protocol (*8*) with a few minor variations and carried out at reduced volume. Briefly, nuclei were sorted into 1 μL lysis buffer (0.19% vol/vol Triton-X 100, 2U SUPERase RNase inhibitor, ERCC RNA spike-ins (ThermoFisher). 1 μL poly-A hybridization mix (final conc. 2.5mM/ea. dNTPs + 2.5 μM oligo-dT primer) was added to each well and the plate was incubated at 72°C 3 min before putting back on ice. 2.85 μL RT reaction mix (final concentration 1μM TSO, 1x Maxima RT buffer (Life

Technologies), 1M betaine, 5 mM DTT, 6 mM $MgCl_2$, 0.5 U SUPERase RNase-inhibitor, 2 U Maxima RT) was added and plate was incubated in a Thermomixer C with ThermoTop (Eppendorf) with the following program:

| | | |
|---|---|---|
| 42°C | 2 min | 2,000 rpm |
| 42°C | 60 min | 1,500 rpm |
| 50°C | 30 min | 1,500 rpm |
| 60°C | 10 min | 1,500 rpm |

or in a thermocycler with the following program: 42°C 90', 10 x [50°C 2', 42°C 2'], 70°C 15'. After the RT reaction, 7.5 μL pre-amp PCR mix (final conc. 1x KAPA HiFi HotStart Readymix (Kapa Biosystems), 0.1 μM IS PCR primer) was added to each well, and plate was incubated in thermocycler: 3' 98°C, [cycle #] x [98°C 20", 67°C 15", 72°C 6'], 72°C 5'. Number of pre-amplification cycles used varied between 18-21, but had little effect on final library quality or complexity. Full-length cDNA was cleaned up using a 0.8x Ampure XP protocol (Beckman Coulter). Final libraries were built from successful cDNA preps using the Nextera XT kit (Illumina) with reduced volume (1/4 or 1/5 standard volumes). Positive control samples from the first part of the protocol were replaced with water (no-DNA controls) before performing Nextera prep. Up to 384 libraries were multiplexed together and sequenced on an Illumina HiSeq 2000 using a 40bp single-end protocol, or on an Illumina NextSeq using a 40x40bp paired-end protocol.

Primer sequences:
oligo-dT:
/5BiosG/AAGCAGTGGTATCAACGCAGAGTACTTTTTTTTTTTTTTTTTTTTTTTTTTTTTTVN
Template-switching oligo (TSO): /5Biosg/AAGCAGTGGTATCAACGCAGAGTACATrGrG+G
IS PCR primer: /5BiosG/AAGCAGTGGTATCAACGCAGAGT

<u>snRNA-seq data processing</u>

Reads were trimmed and quality-filtered using Trim Galore v.0.4.1 (*32*) and aligned using STAR v.2.7.1a (*33*). To minimize mapping bias in favor of the reference strain (Col), reads from Col-Cvi crosses were mapped to a Col-Cvi 'metagenome', consisting of the TAIR10 sequence appended to a Cvi 'pseudogenome' generated by substituting the Cvi allele at 576,697 Col-Cvi SNPs (*17*). Similarly, reads from Col-L*er* crosses were mapped to a Col-L*er* 'metagenome' created using 382,686 L*er* SNPs. Sequences from ThermoFisher ERCC RNA spike-ins were also appended to the metagenome. Reads that mapped uniquely to the ERCC sequences were extracted and omitted from the rest of the analysis. Reads with a single best alignment to the Col-Cvi or Col-L*er* metagenomes or with exactly two equal best alignments, each to equivalent positions on the Col and Cvi/L*er* chromosomes, were considered uniquely mapping. Procedures and scripts for mapping with the metagenome are available in (*34*). Reads overlapping a SNP were identified explicitly using a custom script (assign_to_allele.py, *34*) and assigned to parent-of-origin based on allele. All SNPs within a read had to agree on parent-of-origin for the read to be considered allele-specific. PCR duplicates were removed using MarkDuplicates, from the Picard Toolkit (*35*). Total and allele-specific counts over genes were then obtained using htseq-count v.0.9.1 (*36*) and the Araport11 gene annotations (excluding new Araport11 annotations antisense to existing TAIR10 genes) (*37*). Single-nuclei samples with a total of at least 1,500 genes detected (≥ 1 overlapping read) and 1,000 genes well-detected (≥ 5 overlapping reads) prior to removing PCR duplicates were considered high quality and kept for subsequent analyses. On average, we detected expression from 3,200 genes per 3C endosperm nucleus and 4,200 genes per 6C endosperm nucleus (Fig. S2). All negative controls (no nucleus sorted) lacked reads mapping to *A. thaliana* (Fig. S2). Despite arising from nuclear RNA, few intronic reads were recovered, though somewhat more than for whole-cell bulk RNA-seq (Fig. S3).

Metaplots of coverage over genes and introns

To generate plots in Fig. S3, bigWig coverage files for each library were generated using the deepTools function bamCoverage (*38*, v.3.2.0) with options -bs 1 --normalizeUsing CPM. Coverage over genes and introns was calculated using deepTools computeMatrix in scale-regions mode with options -a 500 -b 500 -m 2000 -bs 50 for genes and -a 200 -b 200 -m 200 -bs 10 for introns. Average profiles for each library were then obtained using deepTools plotProfile with option --perGroup and extracting matrix file with --matrixFile option. Average and s.d. of coverage across all libraries was calculated for each bin in the profile and used to generate plots in Fig. S3.

SC3 clustering and tissue assignment

Initial clustering of the full count matrix was performed using SC3 (*9*); a custom wrapper script used for these analyses (single_cell_cluster_SC3.R) is included in the Github repository. Genes expressed in fewer than 5 nuclei or with fewer than 10 total reads across all nuclei were omitted from this analysis, with a final set of 22,950 genes used for clustering. Counts were converted to CPM using the calculateCPM() function in the R package scater (*39*) before clustering. Optimal number of clusters was estimated using SC3's built-in algorithm. Benchmarking studies have found that SC3 tends to under-cluster (*40*); we therefore sometimes performed additional sub-clustering on clusters that clearly contained additional subgroups (Fig. 1, Figs. S4, S5, S14).

Initial tissue assignments were made based on both the overall % of maternal reads detected for each nucleus (%mat), and a preliminary clustering using tSNE that strongly separated seed coat and endosperm nuclei. tSNE of all nuclei was performed on CPM values using the runTSNE() function in the scater package (*39*), and projected nuclei were clustered using k-means clustering with k = 3. One of these clusters clearly corresponded to seed coat nuclei based on %mat. Nuclei either in that seed coat cluster or with %mat > 85% were preliminarily assigned to seed coat, while those with %mat < 60% were preliminarily assigned to embryo, and all others were assigned to endosperm. Since the SC3 clustering results (Fig. S4) indicated that clusters were highly tissue-dependent, initial tissue assignments were refined based on the SC3 clusters, such that all nuclei in the same cluster were assigned to the tissue assignment of the majority of nuclei. Only 31 nuclei out of 1,437 (2.16%) had their tissue assignments adjusted based on the SC3 clustering results.

At earlier stages of seed development, seeds contain few endosperm-derived 3C and 6C nuclei relative to diploid-derived nuclei (predominantly seed coat), and 3C/6C nuclei become difficult to sort accurately, particularly for very young (2-3 DAP) seeds (Fig. S1). The 3C population is also generally smaller than the 6C population at early timepoints (2-4 DAP), but becomes larger at later timepoints (5 DAP). Due to these factors, seed coat nuclei were obtained at varying rates, ranging from 0% to > 80% per batch/plate, with higher seed coat contamination both at earlier timepoints and when sorting from the smaller 3C peak compared to the 6C peak (Fig. S1).

Once nuclei were assigned to specific tissues, SC3 was used to cluster nuclei from 4 DAP Col x Cvi (CxV) and Cvi x Col (VxC) $F_1$ endosperm and seed coat separately (Fig. 1B, Fig. S5). For CxV endosperm, the 42 nuclei in the last cluster (cluster 10) were re-clustered using SC3 to further resolve cell types. After comparing the results to the whole-dataset SC3 clustering (Fig. S4), we further separated one of these clusters into clusters 12 and 13 manually, based on the fact that these were in two separate clusters in the full SC3 clustering and likely failed to be separated here due to the smaller number of nuclei. For VxC endosperm, initial clustering produced 8 clusters, A-H. Cluster C (n = 30) was re-clustered into clusters 3 and 4, while clusters F-H (n = 208) were not well-resolved and were also re-clustered into clusters 7-11. For CxV seed coat, SC3 produced 6 clusters and no additional sub-clustering was performed. For VxC seed coat, the last cluster in the initial clustering was further subclustered into two clusters.

Plots of average total or allelic expression of genes or gene groups over clusters

Dot plots of average total expression of either individual genes (e.g. Fig. 2A, Fig. S11) or groups of genes (e.g. Fig. 1C, Fig. S4) were made using a custom script (single_cell_RNAseq_plots.R) available in the Github repository, in 'dot' mode. Briefly, the raw count matrix (rows: n genes x m columns: nuclei) was first converted into $log_2$(CPM) values, and for each gene, average $log_2$(CPM>0) was calculated across all nuclei in each cluster to obtain a matrix of n genes x c clusters. Only nuclei with nonzero CPM values (CPM > 0) were used to calculate the average. The fraction of nuclei in each cluster with at least one detected read (CPM > 0) was also calculated. Plots were generated by indicating average $log_2$(CPM>0) using dot color, and fraction of nuclei with at least one detected read in gene as dot size. For plots aggregating groups of genes (e.g. Fig. 1C), average $log_2$(CPM>0) and fraction of zeros were also averaged across all genes in each group to obtain a single value per group.

Plots of average total or allelic expression of individual genes shown in Fig. 3C,E, Fig. S5D, S6, S21, S24, and S25 were made using the same custom script (single_cell_RNAseq_plots.R) in 'lin' mode. For total expression, average $log_2$(CPM>0) and fraction of zeros were obtained as above. For allelic expression, $log_2$(CPM>0) and fraction of zeros were obtained for maternal and paternal counts separately. Paternal counts from each cluster were also fit to the appropriate distribution (NB or ZINB, see 'Modeling maternal and paternal counts' below), and two values were randomly drawn from this distribution and summed for each nucleus in the cluster to simulate a doubling of paternal dosage (dotted blue lines). Plots in Fig. S15 were made using the same script in 'cmp' mode.

Lists of marker genes from seed compartment RNA-seq data obtained by laser capture microdissection (LCM) of seed tissue *(7)* were obtained from Schon et al. 2017 (*27*). All genes that were considered markers for different tissues at different timepoints (n = 181) were discarded. Markers for globular and heart stages (the stages used in this study) were then kept for analysis (n = 1,945).

Identifying differentially expressed genes

Differentially expressed genes between SC3 clusters were identified using DESingle, which performs well with small numbers of cells, as in several of our clusters (*41,42*). To identify genes that were differentially expressed within our dataset, DEsingle was run on all possible pairs of clusters within the 14 CxV endosperm clusters, within the 11 VxC endosperm clusters, between the CxV and VxC endosperm clusters, within the 6 CxV seed coat clusters, within the 8 CxV seed coat clusters, between the CxV and VxC seed coat clusters, and between each endosperm cluster and each seed coat cluster (741 comparisons). Finally, to control for batch effects that may confound the DE analysis, DEsingle was also used to look for DE genes between all batches (batches = 96-well plates, prepared on different days) within the CxV and VxC seed coat and endosperm (8 batches represented in each of CxV endo, CxV seedcoat, VxC endo and VxC seedcoat, 28 * 4 = 112 comparisons).

For each non-batch comparison, all genes that passed stringent significance cutoffs (pval < 0.0001 and abs[$log_2$(fold change)] > 2) were extracted. These were combined across all comparisons to obtain a list of 4,540 genes that were significantly differentially expressed across at least one experimental comparison (cluster, genotype or tissue). This was repeated for all batch comparisons, resulting in 97 genes called as differentially expressed across at least one pair of batches/plates, suggesting that batch effects in our data are minimal. 40 of the batch differentially expressed genes were also differentially expressed in the main analysis and were censored, yielding a final set of 4,500 differentially expressed genes.

Calculating expression enrichment scores and p-values for gene expression enrichment/depletion in particular clusters or across other factors

Gene expression enrichment scores, which reflect the degree to which a gene's expression is enriched/depleted in a specific endosperm or seed coat cluster relative to other clusters, were calculated using a custom script (cluster_gene_expression.R) available in the github repository. This script uses permutation tests to estimate the degree to which a gene is specifically up/downregulated in a cluster, and to calculate a p-value for the significance of this enrichment in each cluster. Briefly, for each gene, $\log_2$(CPM) values for each gene in each nucleus were averaged across all nuclei in each cluster. Cluster labels were then randomly permuted 1000 times (controlling for various factors, see below), and average $\log_2$(CPM) values were calculated using the shuffled cluster labels for each permutation, yielding a background distribution of 1000 values for each gene+cluster combination. Where applicable, we controlled for tissue type (endosperm vs. seed coat), genotype (CxV vs VxC), and wash (yes/no indicating if nucleus was washed during prep, see snRNA-seq library preparation and sequencing) by only permuting cluster labels among nuclei with the same tissue/genotype/wash. The mean and standard deviation of the n = 1000 permuted values was used to calculate a pseudo-Z-score, called the 'enrichment score', reflecting the degree to which the true observed value $x$ for any given gene,cluster combination is extreme relative to the random distribution estimated by permuting the cluster labels:

$$Z = \frac{x - \mu_B}{\sigma_B}$$

where $\mu_B$ and $\sigma_B$ are the mean and standard deviation of the n = 1000 shuffled values, respectively. 'Enrichment score' matrices were clustered using either k-means clustering (Fig. S7) or hierarchical clustering (Fig. 3A,D). The analysis proceeded similarly for calculating enrichment scores over cell cycle phases, with cell cycle phase taking the place of clusters. Similarly, enrichment scores and p-values over tissue/genotype/wash, where applicable, were also calculated by permuting the labels for tissue/genotype/wash across the different samples, and estimating pseudo-Z-scores and p-values as above.

This analysis was performed using either total expression (e.g. Fig. 3A, Fig. S7) or allelic expression (e.g. Fig. 3B). Analyses of total expression were carried out as above. For allelic expression, the analysis above was carried out over the maternal and paternal expression data separately (cluster_gene_expression.R --method separate), and the difference between the maternal and paternal enrichment scores was plotted as a heatmap (Fig. 3B).

To estimate the probability that a gene's expression was enriched or depleted in a particular cluster, a p-value equal to the fraction of times (out of 1000 permutations) that the observed value $x$ was greater than the shuffled mean was also calculated. If this value was less than 0.025, a gene was considered significantly depleted in that cluster; if greater than 0.975, the gene was considered significantly enriched in that cluster.

GO term analysis

The R package 'topGO' was used to identify GO terms significantly enriched among certain groups of DE genes (*43*). Briefly, GO annotations were obtained from plants_mart at plants.ensembl.org using the 'biomaRt' package (*44*). Gene lists of interest were analyzed using the topGO runTest function, with algorithm = 'elim' and statistic = 'fisher'. The background set of genes (gene universe) was the set of 29428 genes with detectable expression in the full snRNA-seq dataset. For each gene list, all significant GO terms (< 0.005) were obtained. The list of all genes associated with each GO-term was obtained using the topGO genesInTerm() function. For plots showing average expression enrichment scores for GO term-associated genes (Fig. S8, S10), enrichment scores for all gene associated with each GO-term were averaged together. A script for performing this analysis, run_topGO.R, is provided in the Github repository.

Cell cycle analysis

To evaluate the positioning of our single-nuclei samples relative to the cell cycle, we performed a modified 'trajectory analysis' using a custom R script (single_cell_trajectory_analysis.R), available in the github repository. A list of 22 genes associated with various phases of the cell cycle, including G0, was manually curated from the literature (*45,46*). The count matrix (n=1,437 nuclei) was first filtered to remove lowly expressed genes, and count values were converted to CPM. Of the 1,437 nuclei, 1,309 (91%) had expression of at least one of the 22 cell cycle marker genes and were retained for further analysis. t-SNE was first performed over CPM values for the 22 cell cycle genes, using the Rtsne package (*47*) with initial_dims = 2 and perplexity = 100. Projected points were then clustered using k-means clustering with k = 6, corresponding approximately to the G0, G1, G1 to S, S, G2 and M phases of the cell cycle.

Dijkstra's algorithm was used to trace a trajectory that was required to pass through the medoid of each k-means cluster. The trajectory was recalculated 200 times, sampling only 50% of the nuclei each time. Each of the 200 trajectories were then combined (*48*) and averaged to build a single smoothed, consensus trajectory through the tSNE plot. Each point in the plot (representing a single nucleus) was then projected onto the consensus trajectory to estimate the positioning of each nucleus along the cell cycle. To identify other genes that vary significantly according to the cell cycle in our dataset, library depth-normalized counts were obtained using the procedure used by edgeR (*49*). A hurdle model, which uses a truncated distribution to model the additional zeros present in single-cell RNA-seq data and has been used previously in similar contexts (*50*) was used to model the normalized counts. The distribution used for the non-zero component of the hurdle model was the negative binomial distribution, which is used frequently to model count data from RNA-seq (*49,51*). We used this count model to test each gene using a regression approach (hurdle() function in R pscl package, (*52*) to determine whether the cell cycle trajectory explains a significant amount of the variation in each gene's expression. P-values were adjusted using the Bonferroni method, and genes with adjusted *p*-values < 0.01 were considered significantly cell-cycle dependent. This analysis identified a total of 1,065 cell-cycle-dependent genes.

Identifying imprinted genes from snRNA-seq data

Assessing imprinting using snRNA-seq data is complicated by several factors, including dropouts (genes not detected in a cell due to low input & technical factors) and transcriptional bursting kinetics, which can cause transcription at a locus to appear monoallelic at the moment of cell/nucleus capture even if a gene is biallelically expressed (*53-55*). As a result, imprinting must be assessed by aggregating information from multiple single nuclei across the dataset. Additionally, in most angiosperms including Arabidopsis, endosperm has a maternal:paternal (m:p) genome dosage of 2m:1p rather than 1m:1p. mRNAs from the two maternal alleles are indistinguishable, and thus cannot be modeled independently or directly compared to paternal expression, as in existing methods for detecting imprinting from scRNA-seq (*56,57*). We therefore developed a method for assessing imprinting that accounts for maternal and paternal dosage in endosperm (single_cell_ASE_analysis.R, in github repository).

1. Modeling maternal and paternal counts and testing for parental bias

Paternal counts were fit to the negative binomial (NB) and zero-inflated negative binomial (ZINB) distributions, and the Akaike Information Criterion was used determine which had the best fit (Fig. S15). The NB distribution is commonly used to model counts in bulk RNA-seq (*49,51*), while the ZINB extends the NB with an additional parameter to model dropout events and has been used for scRNA-seq analysis (*41*,*58*). To account for maternal genome dosage, maternal count data were instead modeled as the sum of two independent, identically distributed ZINB or NB random variables. The NB has parameters $\mu, \sigma$, representing the mean and variance respectively, and the ZINB has an additional $\nu$ parameter modeling zero inflation (*59*):

| Distribution | Equation | Mean | Variance |
|---|---|---|---|
| $X \sim NB(\mu, \sigma)$ | $P_X(x\mid \mu, \sigma) = \frac{\Gamma(x+\frac{1}{\sigma})}{\Gamma(\frac{1}{\sigma})\Gamma(x+1)}\left(\frac{\sigma\mu}{1+\sigma\mu}\right)^x\left(\frac{1}{1+\sigma\mu}\right)^{\frac{1}{\sigma}}$ | $E[X] = \mu$ | $Var(X) = \mu + \sigma\mu^2$ |
| $X \sim ZINB(\nu, \mu, \sigma)$ | $P(x\mid\nu, \mu, \sigma) = \begin{cases} \nu + (1-\nu) * p_{X'}(0\mid \mu, \sigma) & if\ x = 0 \\ (1-\nu) * p_{X'}(x\mid \mu, \sigma) & if\ x = 1,2,3 \ldots \end{cases}$ where $X' \sim NB(\mu, \sigma)$ | $E[X] = (1-\nu)\mu$ | $Var(X) = \mu(1-\nu) * (1 + (\nu + \sigma)\mu)$ |

Parameter estimates were obtained by Maximum Likelihood (*60*). Maternal and paternal parameter estimates were well-correlated with each other (Fig. S16), suggesting that the overall transcriptional kinetics of the maternal and paternal alleles are similar for most genes in our dataset. Both imaging studies and single-cell studies have found that the two alleles of most genes function independently and share similar kinetics (*56*).

Our analysis revealed minor but consistent skew in favor of higher maternal allele expression at most genes, and maternal (1-$\nu$) and $\mu$ estimates were consistently slightly higher than paternal estimates (Fig. S16). This effect was more pronounced in Col x Cvi than in Cvi x Col $F_1$ nuclei, suggesting that technical factors, such as mapping bias in favor of the sequenced strain Col, played a role, though we cannot rule out biological factors (also see 'identifying imprinted genes below'). In light of this, we tested an adjusted null hypothesis for deviation from the expected 2m:1p ratio that accounted for the average skew in the data.

Under the null hypothesis that the gene was not imprinted and that the maternal and paternal ZINB/NB parameters were equal after adjusting for average maternal skew, p-values were obtained using a likelihood ratio test.

2. Evaluating our approach using simulated count data

To test the power and accuracy of our approach, we simulated maternal and paternal count data resembling the CxV and VxC datasets, but with varying levels of overall expression, parental bias, nuclei-to-nuclei variability, and number of nuclei (Fig. S17). Counts were drawn from ZINB distributions with known parameters. Our approach was able to accurately identify genes with parental bias under a wide range of conditions, with accuracy increasing as expression levels or parental bias increased. Simulations included a degree of maternal skew similar to the one in the real data, and showed that failing to adjust for this skew when testing for parental bias led to high false positive maternal bias calls, while an adjusted hypothesis had very few false positives (Fig. S17).

Our method also identified maternal and paternal bias in simulations of highly expressed or highly biased genes from as few as 5 or 10 simulated nuclei, though with a high false negative rate (Fig. S17). Our approach therefore likely has a high false negative rate for lowly expressed genes or genes expressed only in rare subpopulations. For example, *SUVH7* and *MEA* are both well-known imprinted genes (*61,62*), but were not among the PEGs and MEGs identified in this study. However, *SUVH7* expression is limited to the chalazal cyst, a relatively rare population in our dataset. Although our method found that *SUVH7* is clearly paternally biased in VxC, which has more cyst nuclei, we lacked power to identify bias in CxV, which has only 6 cyst nuclei. Similarly, *MEA* displays maternal bias in the data, but is only strongly expressed in CxV E14, a cluster with only 11 nuclei. Therefore, our analysis likely undercounts the true number of imprinted genes. However, false positive rates were low across all conditions tested, suggesting that our method has high specificity but tends to be conservative (Fig. S17).

3. Identifying imprinted genes

Using the method outlined above, *p*-values for significant parental bias were generated for each gene in that had at least 10 allelic counts, and corrected using the Benjamini-Hochberg method (*63*). The 4 DAP CxV and VxC datasets were analyzed separately. Genes with adjusted *p*-value $< 0.05$ and a $\log_2(m/p)$ value more than 0.5 away from the median $\log_2(m/p)$ value across all genes were considered significantly biased (Fig. S18). Genes were divided into weak, medium, or strong bias based on $\log_2(m/p)$ values, and results from both CxV and VxC datasets were combined into a final 'overall status': MEGs/PEGs (maternal/paternal bias in both crosses), Col/Cvi bias (bias in favor of the same strain in both crosses), bias in only one cross, or no bias (Fig. S18, Fig. S19). Unbiased genes with fewer than 20 allelic counts overall likely lacked statistical power and were classified as having 'too few reads'.

Of the MEGs and PEGs previously identified from whole-endosperm RNA-seq (17) with sufficient data for our analysis, most were also considered imprinted based on our dataset (Fig. S20). However, a number of the MEGs and PEGs identified in this study were not previously identified (Fig. S20). Some of these lacked data in the previous study, either because they were not detected or the annotation did not yet exist. However, a number of the MEGs identified in this study were censored from the analysis in the previous study because laser-capture microdissection datasets suggest that these genes are more highly expressed in the seed coat than in the endosperm (*7*). An ongoing concern with bulk endosperm datasets is that contamination from the seed coat, a maternal tissue, can lead to false positives among MEGs; this type of contamination has been shown to occur in some manually dissected endosperm datasets (*27*). However, with our data, it is possible to carefully separate nuclei derived from seed coat from nuclei derived from endosperm based on a number of expression features (Fig. 1), so our CxV and VxC endosperm datasets should not have seed coat contamination. Thus, many MEGs identified here are indeed more highly expressed in seed coat than in endosperm (Fig. S20, S21, *7*). After also noting a widespread skew towards maternal expression in the data (Fig. S16, S19), we tested the possibility that this phenomenon was caused by mRNAs from the seed coat being carried over in the sorting buffer during FANS by adding an additional wash step prior to sorting (see 'Seed Nuclei FANS' above). Surprisingly, endosperm nuclei that had been treated with the extra wash step showed an even more pronounced maternal skew compared to unwashed nuclei, and washed nuclei had even stronger maternal expression of most MEGs compared to unwashed nuclei (Fig. S22). It remains unclear whether this phenomenon is biological, or a consequence of technical factors related to snRNA-seq data.

However, overall these results suggest that our approach is able to identify imprinted genes that could not previously be evaluated by bulk RNA-seq.

**Supplementary Text**

Defining micropylar and peripheral endosperm nuclei cluster identity

After fertilization, Arabidopsis endosperm undergoes three rounds of rapid synchronous nuclear replication without cytokinesis, forming a syncytium of nuclei surrounded by cytoplasm (*3,4*). Then, three morphologically distinct domains are formed: the micropylar, peripheral, and chalazal endosperm regions (Fig. 1E, *5*). Nuclei divide synchronously within a domain but asynchronously with other domains. At one pole of the seed, the dense micropylar endosperm surrounds the embryo, while at the other pole the chalazal endosperm lies adjacent to the termination of maternal vascular tissue. The peripheral endosperm lies between the poles and consists of a large central vacuole with nuclei arranged along the outer periphery, connected by cytoplasmic bridges (*4*). Several days post-fertilization, endosperm cellularization proceeds in a wave from the micropylar to chalazal pole. The chalazal endosperm does not fully cellularize.

We first evaluated the expression of previously defined marker genes for micropylar, peripheral, and chalazal endosperm tissue obtained from published microarray analysis of laser capture microdissection (LCM) of wild-type Ws-0 seeds at a similar stage of development (*7, 27*). Markers for each endosperm domain were enriched in several clusters, while some clusters lacked enrichment for markers of any domain (Fig. 1C). To further refine cluster identity, we identified differentially expressed genes by performing all possible comparisons among nuclei clusters (Fig. S7). GO term analysis indicated that several probable peripheral and micropylar endosperm clusters had increased expression of genes characteristic of S- and M-phase (Fig. 1D, Fig. S10), suggesting that some of the variability between endosperm nuclei could be attributed to differences in cell cycle stage. We therefore performed a trajectory analysis to map each nucleus onto a linear path representing progression through the cell cycle (Fig. S12) (*45,46*). To determine the extent to which cell cycle differences were driving endosperm nuclei clustering, we repeated the clustering while omitting 1,065 genes whose expression varied significantly along the cell cycle trajectory. Similar endosperm clusters were recovered, indicating that cell cycle stage is not sufficient to explain the observed clustering (Fig. S14).

Multiple endosperm nuclei clusters showed elevated expression of genes related to M-phase, such as mitotic chromosome condensation and cytokinesis (Fig. 1D, Fig. S10). To examine where these nuclei were distributed in the endosperm, we performed RNA *in situ* hybridization for AT4G11080/*3XHMG-BOX1*, which was most frequently expressed in VxC E2 and CxV E4, E8 and E10 nuclei (Fig. 2A, Fig. S11). In each of these clusters, the majority of nuclei were identified as being in M-phase (Fig. S12). Hybridization signal was detected in a spotty pattern in the embryo, in the micropylar endosperm, and along the cellularization boundary of the peripheral endosperm (Fig. 2B, Fig. S11), consistent with dividing nuclei. Among the clusters expressing AT4G11080, VxC E2 and CxV E4 had the strongest expression of other M-phase related genes (Fig. S7), had similar overall expression patterns (Fig. S9), and were enriched in genes associated with peripheral endosperm (Fig. 1C). Based on these distinct lines of evidence, we concluded that both VxC E2 and CxV E4 clusters represent M-phase peripheral endosperm nuclei (Fig. 1E). CxV E8, consisting of only 6 nuclei, appears to be a distinct class of possibly mid M-phase peripheral endosperm nuclei (Fig. 1E).

In contrast, CxV E10 and VxC E4 showed some upregulation of M-phase related genes (Fig. 1D) and AT4G11080/*3XHMG-BOX1*, but were transcriptionally distinct from the peripheral M-phase clusters (Fig. S7) and had some expression of LCM micropylar endosperm markers (Fig. 1C). Yet only one cluster, VxC E3, was strongly enriched for the expression of LCM micropylar endosperm markers (Fig. 1C, Fig. S9). However, VxC E3 shared some gene expression features with VxC E4 (Fig. 1B, Fig. S7) and CxV E9, E10, and E11, including expression of AT1G09380/*UMAMIT25*, which encodes an amino acid exporter (*64*) (Fig. 2A). We found that *UMAMIT25* transcript was detected in the micropylar endosperm in both crosses and in 25% of Col x Cvi seeds it was also detected on the leading edge of the cellularization boundary in peripheral endosperm (Fig. 2B). RNA was also detected in the seed coat, as expected based on snRNA-seq data from the seed coat clusters (Fig. S11). This suggests that both VxC E3 and E4 represent distinct populations of micropylar nuclei. VxC E4 and CxV E10 both express M-phase related genes (Fig. 1D) and have similar expression patterns (Fig. S7), suggesting that together CxV E10 and VxC E4 correspond to actively dividing micropylar nuclei. CxV E11 has upregulation of S-phase related genes (Fig. 1D) and may correspond to micropylar nuclei in S-phase. Unlike these clusters, cell cycle trajectory analysis indicated that VxC E3 contains nuclei only in G2 or G0, suggesting these are fully differentiated, non-dividing micropylar nuclei (Fig. 1E, Fig. S12). In Col x Cvi seeds, *RGE1*/*ZOU*/AT1G49770, which is expressed in the embryo surrounding region of the micropylar endosperm and is required for embryo cuticle formation (*65*), is highly expressed in CxV E9, but not E10 or E11. This suggests that CxV E9 represents the micropylar nuclei in the embryo surrounding region in CxV (Fig. 1E).

Several clusters corresponded to peripheral endosperm, which comprises the bulk of the nuclei in endosperm (Fig. 1D, E). While some were most strongly associated with a particular cell cycle phase, a distinctive subset were characterized by upregulation of the photosynthetic machinery and were classified as ‘energy-generating’ peripheral endosperm. These three clusters were distinguished from each other by GO terms specific to certain clusters, such as translation and fatty acid biosynthesis for energy-generating peripheral I (Fig. 1D, Fig. S10). This suggests that there are distinct nuclei populations within peripheral endosperm whose function is primarily to carry out photosynthesis and biosynthesis of different nutrients. In contrast to these clusters, VxC E5 was enriched for down-regulation of photosynthesis genes and fructose 1,6 bisphosphate metabolism, but had few upregulated genes overall and appeared relatively metabolically inactive. This cluster was therefore classified as ‘inactive’ peripheral endosperm (Fig. 1E).

<u>Defining seed coat nuclei cluster identity</u>

The developing seed coat consists of five distinct cell layers and the chalazal seed coat region (Figs. S5,S6). The innermost cell layer ii1, or endothelium, is characterized by elevated production of proanthocyanidins (*66*). Genes related to proanthocyanidin production were strongly upregulated in VxC E7 and E8 (Fig. S5). Two other clusters, CxV S4 and VxC S4, had high expression of *BAN*, a gene known to be expressed in the endothelium, but not *TT12*, another endothelium marker. By contrast, CxV S6 and VxC S7, S8 had high expression of both genes (Fig. S6). Based on these and other gene expression data, we characterized CxV S4 and VxC S4 as ‘potential endothelium’ and VxC S7, S8 as ‘proanthocyanidin-synthesizing endothelium’. The ‘potential endothelium’ clusters were also characterized by enriched expression of genes involved in dNTP synthesis and glutamine metabolism (Fig. S5). Clusters CxV S2 and VxC S2 likely correspond to the epidermal layer (oi2). Both clusters expressed the epidermal cell fate transcription factor *GLABRA2*, and were enriched for cutin biosynthesis and transport (Fig. S5). The clusters with the most nuclei, CxV S1 and VxC S1, had elevated expression of genes related to the synthesis of flavonol, which is produced in the outer integument layers (Fig. S5, *67*). The chalazal seed coat clusters, CxV S3 and VxC S6, were enriched for sucrose biosynthesis, mannose response, and glucose response, all consistent with a nutrient transfer function for this region. The chalazal seed coat also had elevated expression of multiple transcription factors, including the MADs domain transcription factors

*AGL15*, *AGL16*, *AGL18*, *AGL2/SEP1*, *AGL69/MAF4*, and the interacting factors *AGL9/SEP3* and *AGL11/STK*, which influence mechanical properties of the seed coat (*68*).

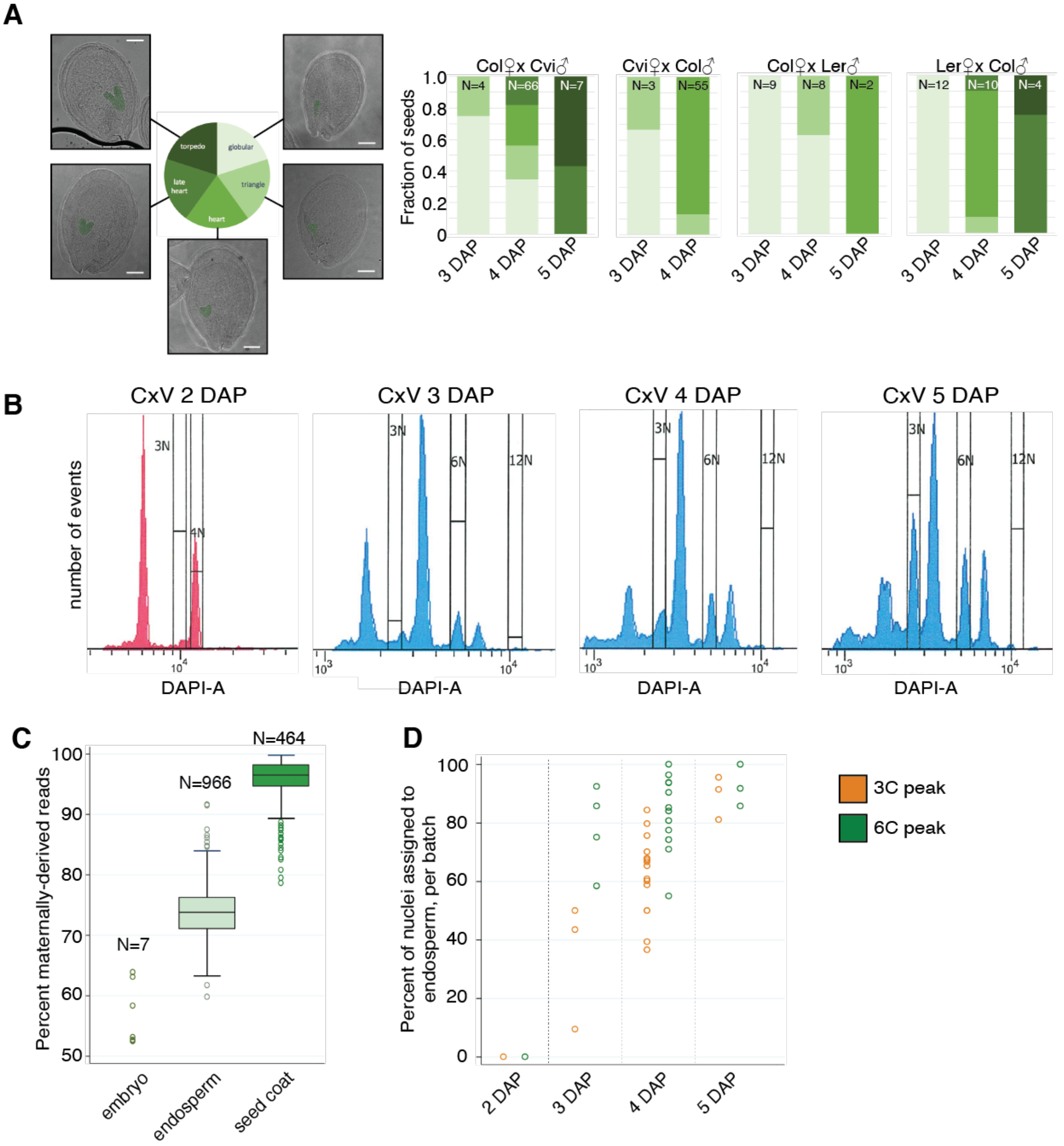


**Fig S1. Seed developmental stages assayed, FANS profiles, and impact on endosperm enrichment.** (A) Summary of seed developmental stages in the different genotypes and timepoints assayed. Number of seeds imaged for each bar shown at top. (B) FANS sorting profiles of Col x Cvi (CxV) seeds at 2 DAP (sorted 09/26/17), 3 DAP (08/10/17), 4 DAP (11/16/17) and 5 DAP (11/14/17). The 2 DAP sample was processed on a different FACS machine than the other three samples. (C) Percent of allelic reads that were derived from the maternally inherited allele, for nuclei assigned as embryo, endosperm, and seed coat (see methods). (D) Percent of nuclei per batch (96-well plate) assigned to endosperm. Nuclei from later timepoints, as well as from the 6C peak, are more likely to correspond to endosperm than nuclei from earlier timepoints or from the 3C peak.

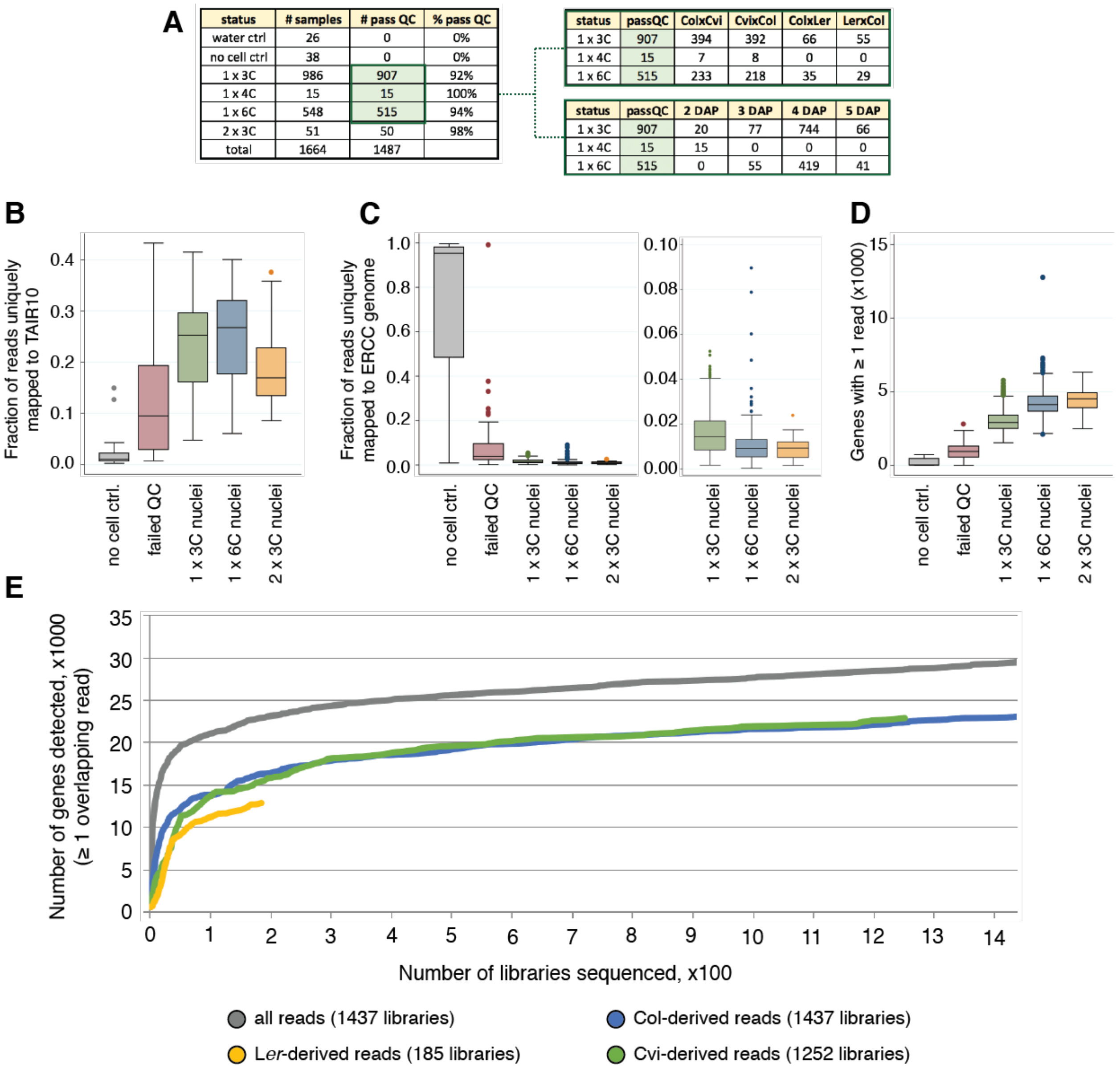

| status | # samples | # pass QC | % pass QC |
| --- | --- | --- | --- |
| water ctrl | 26 | 0 | 0% |
| no cell ctrl | 38 | 0 | 0% |
| 1 x 3C | 986 | 907 | 92% |
| 1 x 4C | 15 | 15 | 100% |
| 1 x 6C | 548 | 515 | 94% |
| 2 x 3C | 51 | 50 | 98% |
| total | 1664 | 1487 | |

| status | passQC | ColxCvi | CvixCol | ColxLer | LerxCol |
| --- | --- | --- | --- | --- | --- |
| 1 x 3C | 907 | 394 | 392 | 66 | 55 |
| 1 x 4C | 15 | 7 | 8 | 0 | 0 |
| 1 x 6C | 515 | 233 | 218 | 35 | 29 |

| status | passQC | 2 DAP | 3 DAP | 4 DAP | 5 DAP |
| --- | --- | --- | --- | --- | --- |
| 1 x 3C | 907 | 20 | 77 | 744 | 66 |
| 1 x 4C | 15 | 15 | 0 | 0 | 0 |
| 1 x 6C | 515 | 0 | 55 | 419 | 41 |



**Fig S2. Summary of library quality and comparison to controls.** (A) Left: summary of libraries obtained: water ctrl = Nextera XT library built with water (no DNA); no cell ctrl = no nucleus sorted into well, but carried through the entire prep; 1 x 3C = single nucleus from 3C FANS peak, 1 x 4C = single nucleus from 4C FANS peak (2 DAP only, due to no visible 3C peak); 1 x 6C = single nucleus from 6C FANS peak, 2 x 3C = two nuclei sorted from 3C peak. The 1,437 non-control libraries highlighted in green passed basic QC cutoffs (see methods). Right: 1,437 high-quality libraries by ploidy and genotype (top) or stage (bottom). (B-D) Distribution in control, failed, and high-quality libraries, of (B) the fraction of reads that mapped to TAIR10, (C) the fraction of reads that mapped to the ERCC spike-in genome, and (D) the number of genes detected. 1x3C, 1x6C and 2x3C are libraries that passed QC filters. In (C), panel on right is zoomed-in view of libraries that passed QC. (E) Total number of genes detected as the number of single nuclei sequenced that passed QC increases, defined as at least 1 read mapping to gene. For allelic expression, gene detection was defined as at least 1 allele-specific read (read overlapping a SNP) mapping to gene.

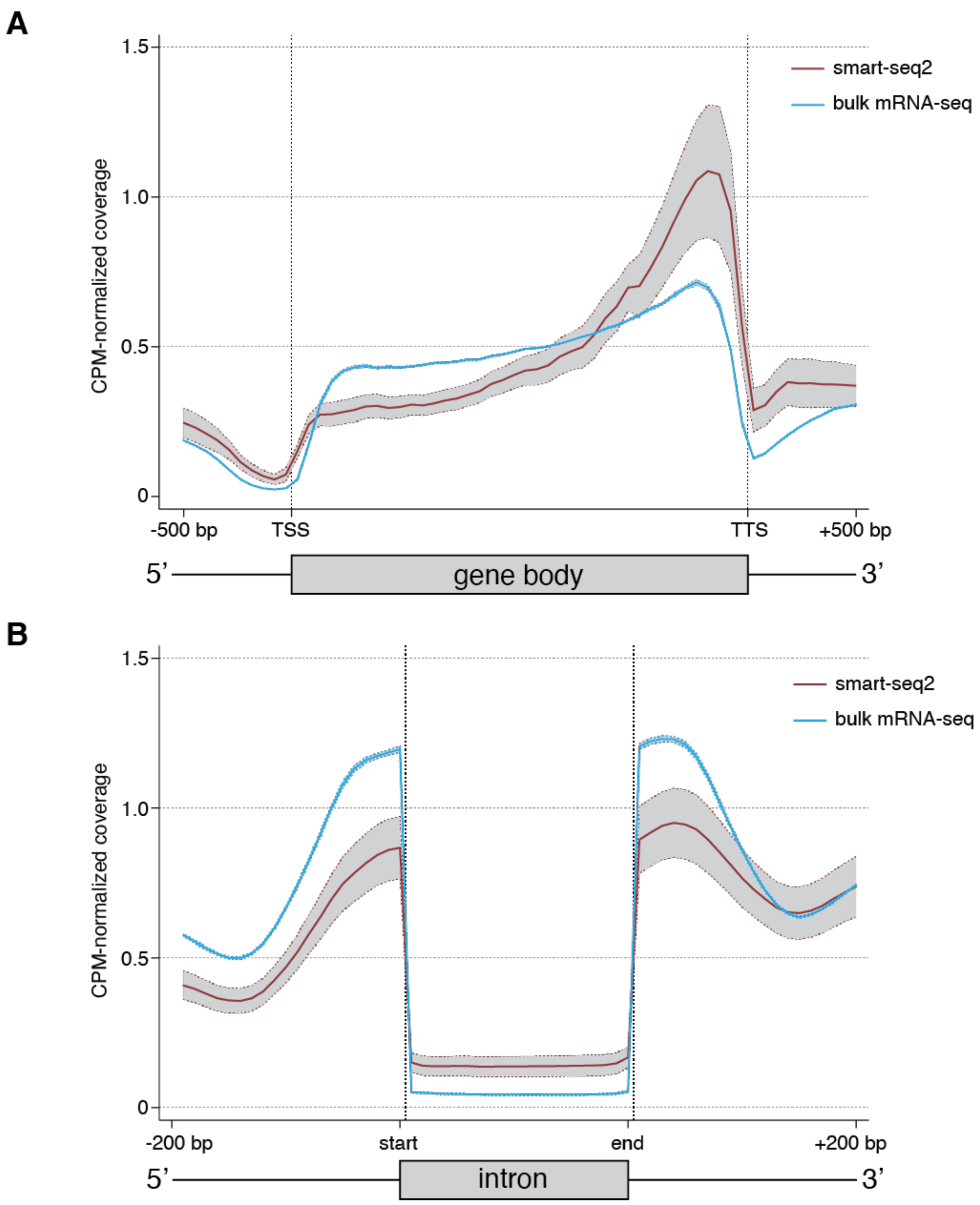


**Fig. S3. Comparison of bias in gene and intron coverage in smart-seq2 libraries vs. published bulk mRNA-seq data.** Single-nuclei smart-seq2 libraries generated in this study were compared to 6 published bulk whole-cell RNA-seq libraries (3 replicates each of Col and Cvi parent lines, *69*). Central solid line indicates overall average, red for Smart-seq2 libraries (average of all 1437 Smart-seq2 libraries that passed QC) and blue for bulk mRNA-seq (average of 6 libraries, *69*). Dotted lines and shaded area indicate +/- 1 SD.

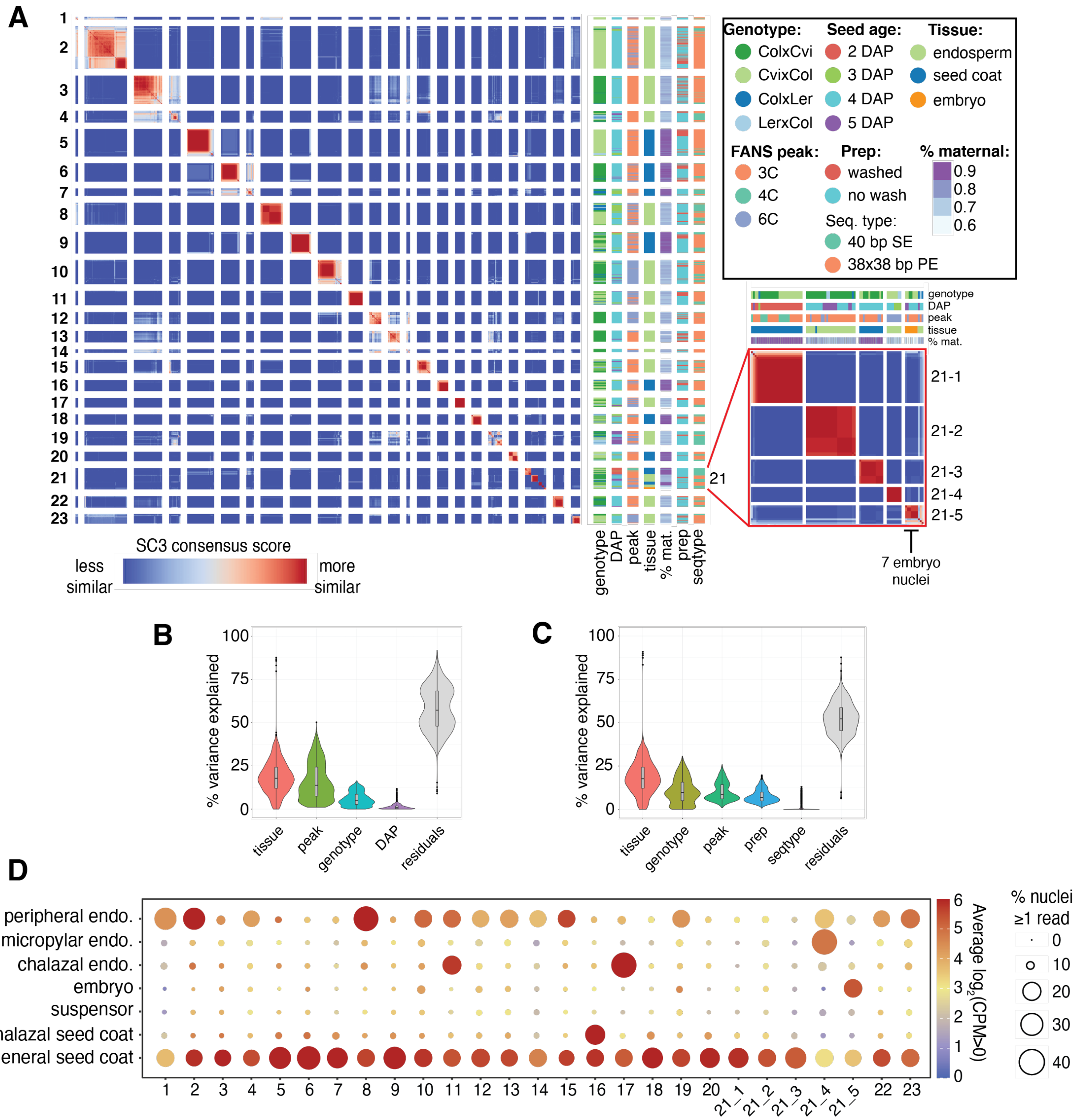


**Fig S4. Clustering of all 1437 high-quality nuclei in the dataset.** (A) Heatmap of SC3 clustering of all 1437 nuclei. Genotype, FANS peak, prep method (see 'Seed nuclei FANS'), sequencing type, % maternal (percent of allelic reads derived from maternal allele), and seed age also shown. (B) Partitioning of the variance in CPM values for the 1437 nuclei samples, according to tissue, peak, genotype and DAP, using the R package 'variancePartition' (*70*). (C) Same as (B), over the 1096 Col x Cvi and Cvi x Col 4 DAP samples only. In this group, prep and sequencing type are less confounded with sources of biological variation (e.g. all washed samples are either Col x Cvi or Cvi x Col 4 DAP, so prep is confounded with genotype and DAP in the full dataset), so their contribution to the variation could be more reliably estimated. (D) Average expression of marker genes for various seed compartments (globular and heart stage) *(7,27)* for nuclei in each cluster. Size indicates the average percent of nuclei with >0 counts, color indicates average $\log_2$(CPM) for all nuclei with CPM > 0.

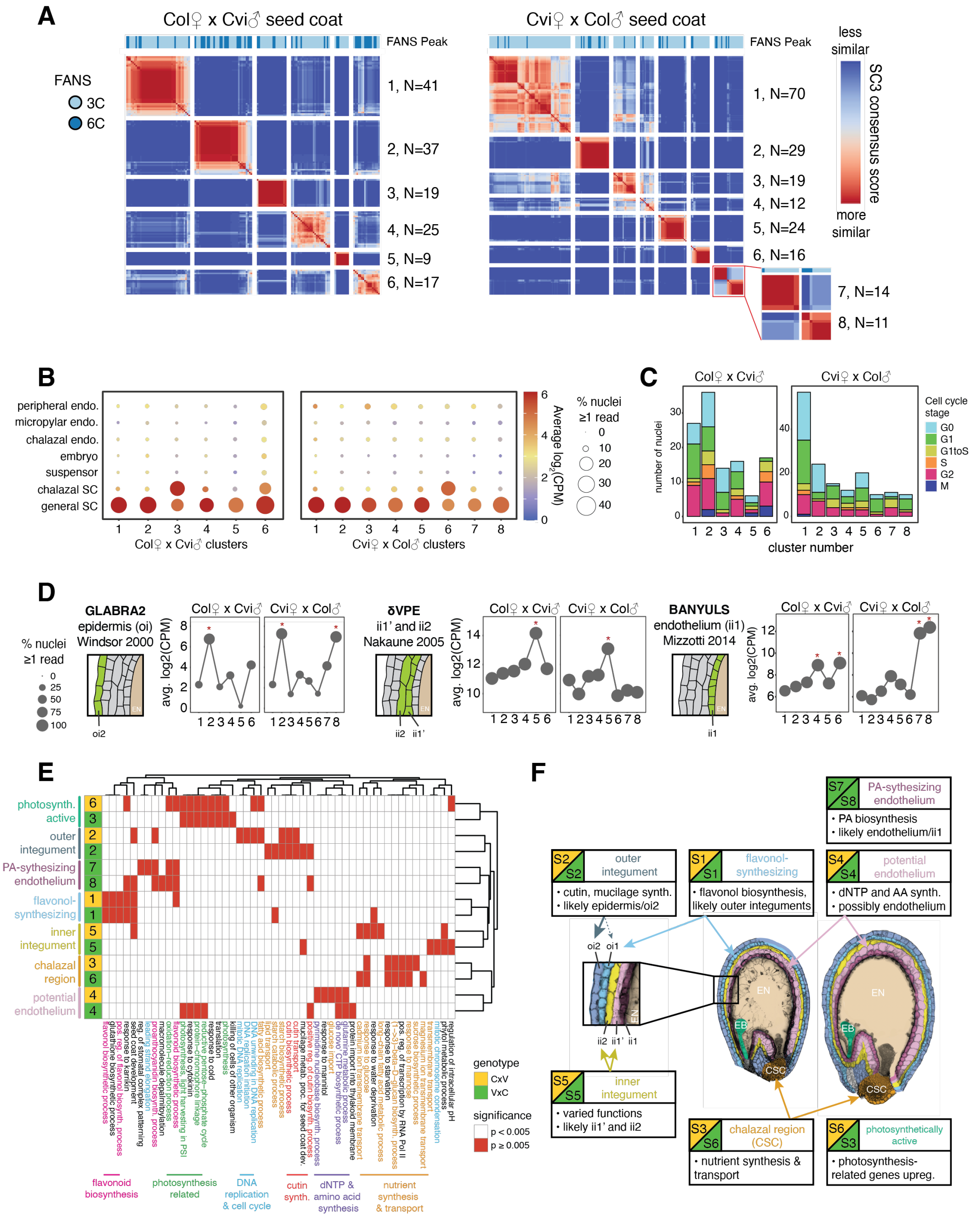


**Fig S5. Characterization of seed coat nuclei.** (A) SC3 clustering of 4 DAP seed coat nuclei. (B) Average expression of LCM seed tissue markers (*7, 27*), over seed coat clusters. Dot color: average $log_2$(CPM); dot size: average percent nuclei with CPM>0. (C) Cell cycle phase by cluster. (D) Average expression of genes specific to particular seed coat cell layers (*71-73*) across nuclei clusters. Schematic of seed coat cell layers, from ii1 (the endothelium, innermost) to oi2 (epidermis, outermost); layers where expression was observed in indicated study highlighted green. Red star: significantly higher expression in cluster (permutation test, $p < 0.05$). (E) Top 5 GO terms for significantly upregulated genes in each cluster. (G) Cluster identities and characteristics; false-colored Col x Cvi (left) and Cvi x Col (right) seed images. EB = embryo, EN = endosperm, CSC = chalazal seed coat. Inset: the five seed coat cell layers.

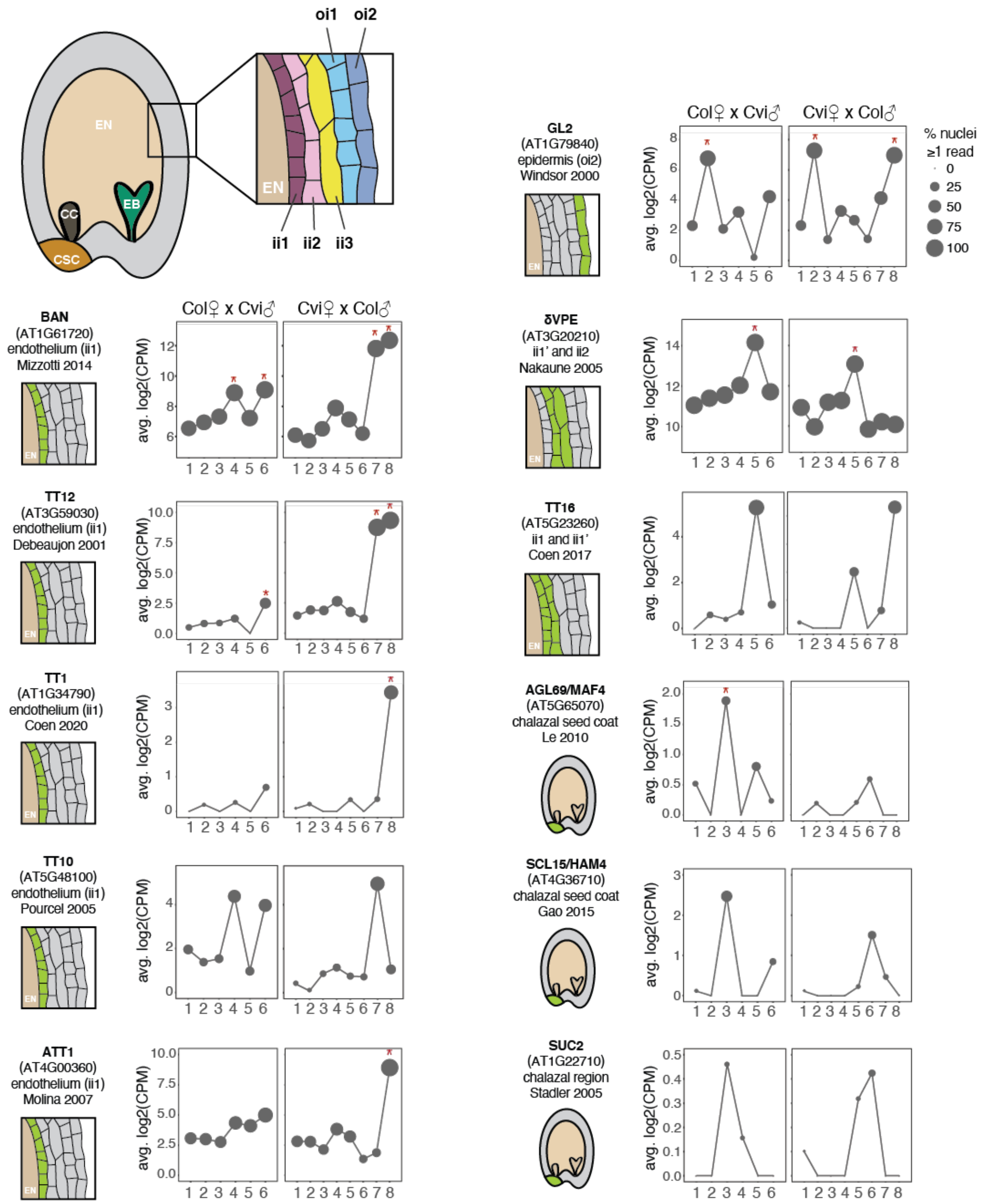


**Fig S6. Expression of genes with known cell layer or region-specific seed coat expression patterns in seed coat nuclei clusters.** Genes were curated from the literature and only considered if expression pattern was determined either by GUS staining or *in situ* hybridization (*67,71-79*) Schematic at left of each plot shows the localization of the gene according to the indicated study. Red star: gene was significantly upregulated in that cluster ($p < 0.05$, permutation test). TT1, TT10, TT16, SCL15 and SUC2 were too lowly expressed to evaluate statistical significance. (Top Left) Schematic of seed showing location of the chalazal seed coat region (CSC), as well as embryo (EB), endosperm (EN), and chalazal cyst (CC). Inset: schematic of the five cell layers in seed coat, from ii1 (the endothelium, innermost cell layer) to oi2 (epidermis, outermost cell layer).

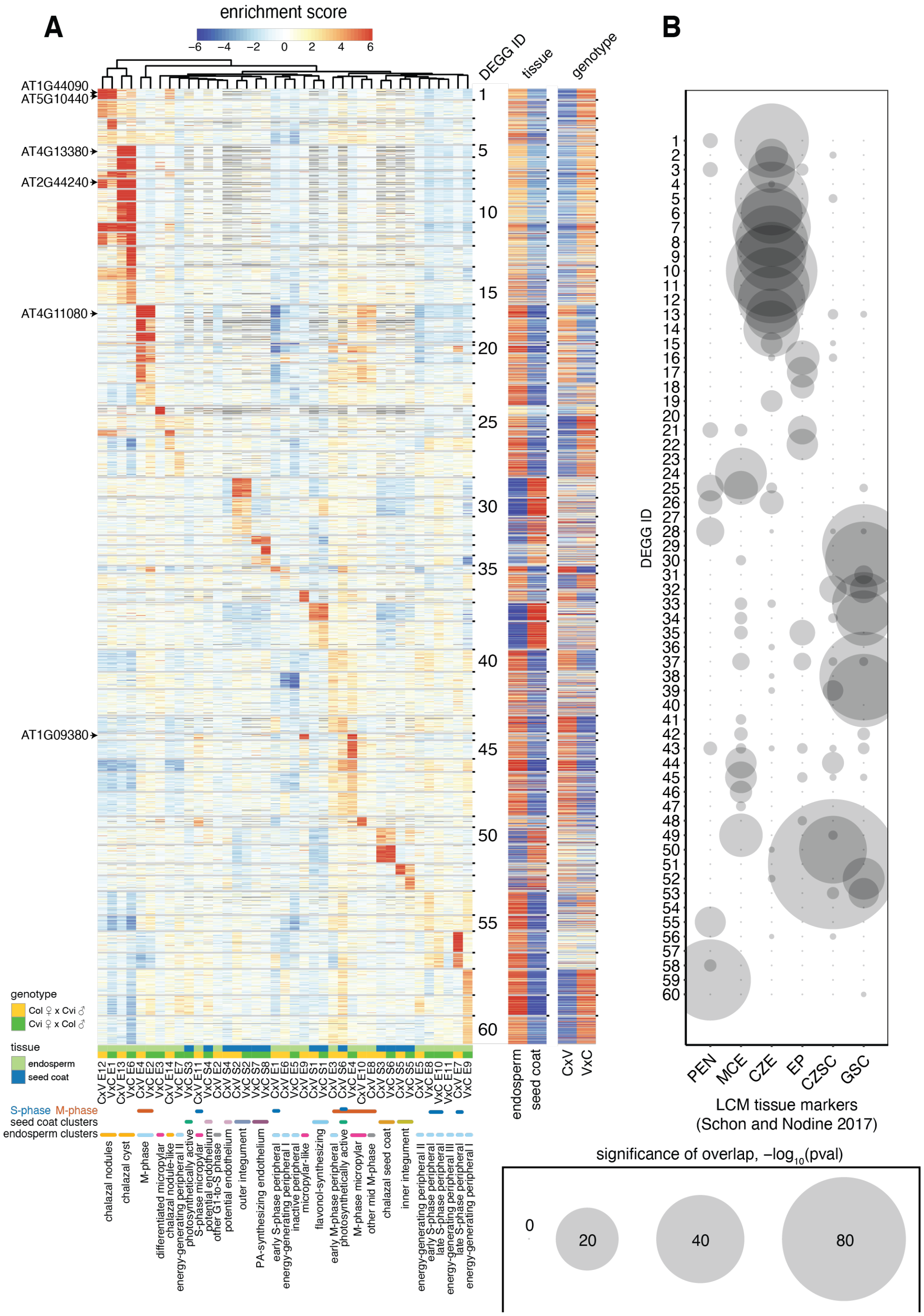


**Fig. S7. Overall expression patterns of 4,500 genes significantly differentially expressed among the endosperm and seed coat clusters.** (A) Heatmap of gene expression 'enrichment scores' for all 4,500 genes clustered by k-means clustering (k =60). Genes used for *in situ* hybridization (Fig. 2, Fig. S11) indicated at left. Nuclei clusters are labeled according to cluster identity (Fig. 1E); clusters enriched for genes related to M-phase or S-phase (Fig. 1D) also indicated. Columns on right indicate relative enrichment score in seed coat vs. endosperm and Col x Cvi vs. Cvi x Col. (B) Significance of overlap between genes in each k-means cluster from (A), and marker genes for different seed compartments *(7,* curated by *27*). P-values were computed using the hypergeometric test.

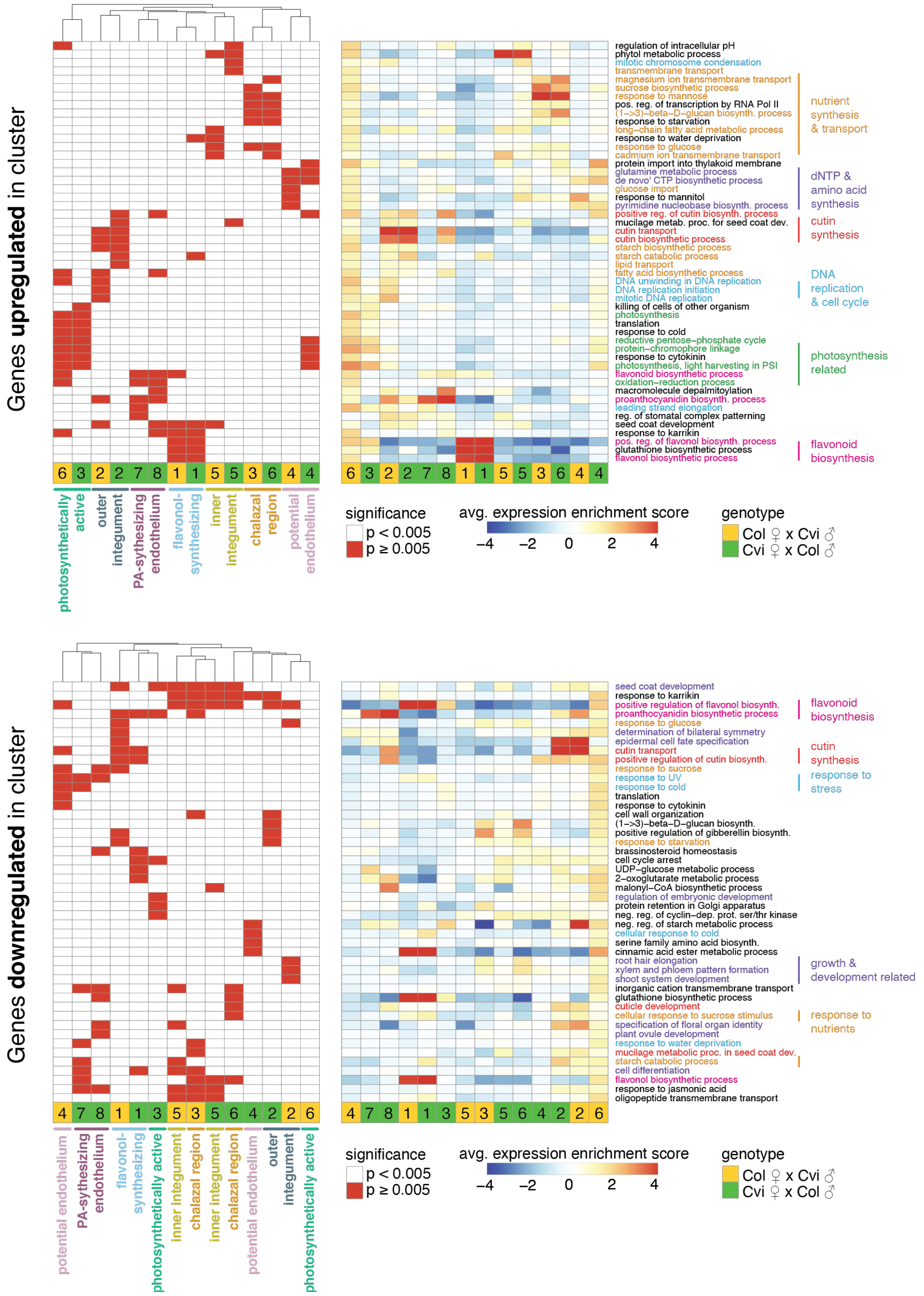


**Fig. S8. Heatmaps of the 5 most significantly enriched GO terms among genes upregulated (top) and downregulated (bottom) in each seed coat cluster.** Significant terms are flagged in left heatmap, while average expression 'enrichment score' across all genes associated with GO term is shown at right. Average includes any genes associated with the GO-term that are not significantly up/downregulated in the indicated cluster, so average may not reflect expectations. Order of rows/columns same for left and right heatmaps.

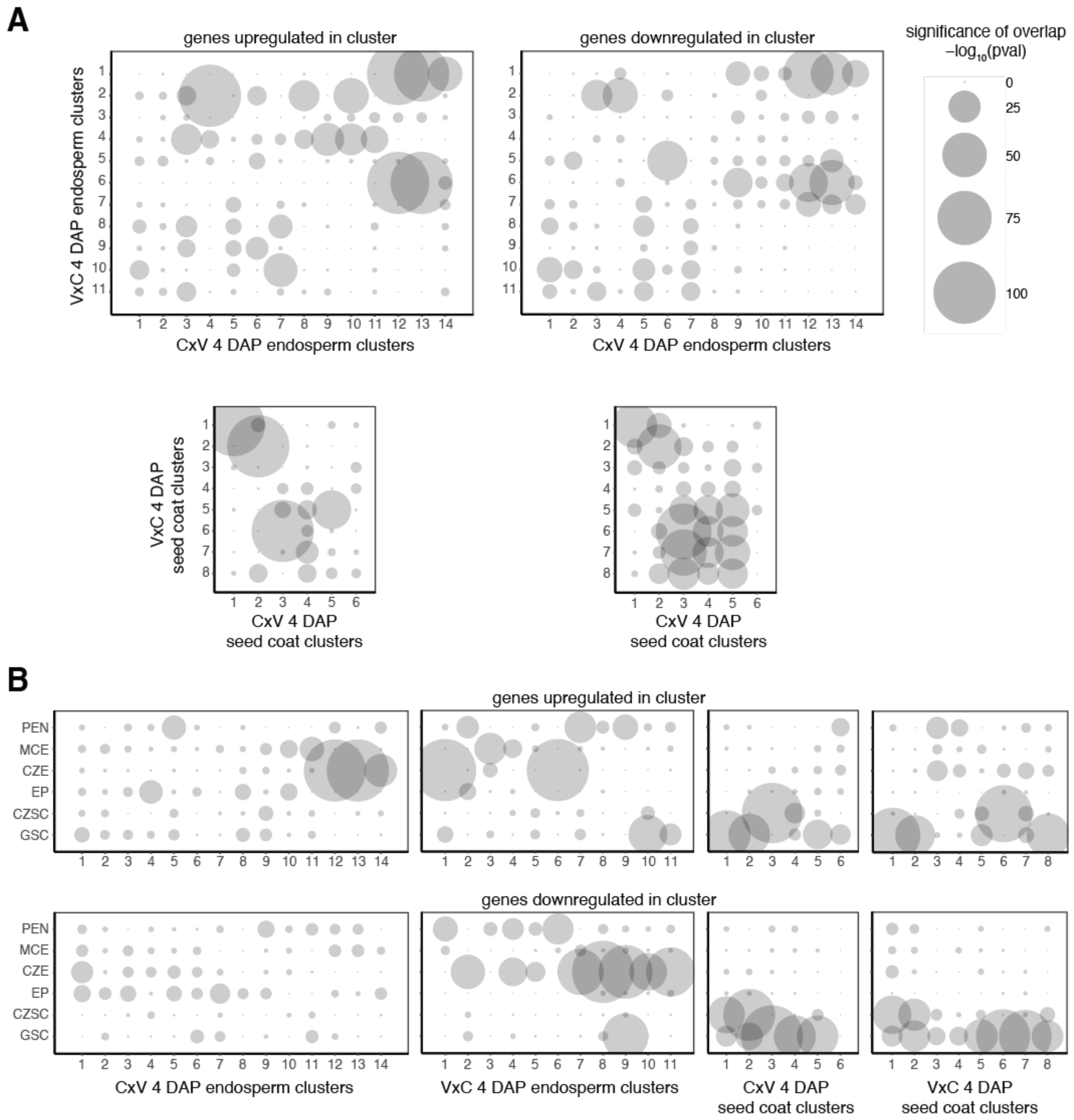


**Fig. S9. Overlap between significantly up- and down-regulated genes across all 4 DAP nuclei clusters and with published seed domain markers.** (A) Significance of overlap of significantly up- and down-regulated genes between Col x Cvi and Cvi x Col 4 DAP endosperm or seed coat clusters. p-value obtained using hypergeometric test. Values above 100 (-log(pval)) were truncated to 100 for plotting. (B) Significance of overlap between genes significantly up- or down-regulated in the nuclei clusters, and marker genes for six seed tissues (*7, 27*). Legend same as A. PEN = peripheral endosperm, MCE = micropylar endosperm, CZE = chalazal endosperm, EP = embryo proper, CZSC = chalazal seed coat, GSC = general seed coat

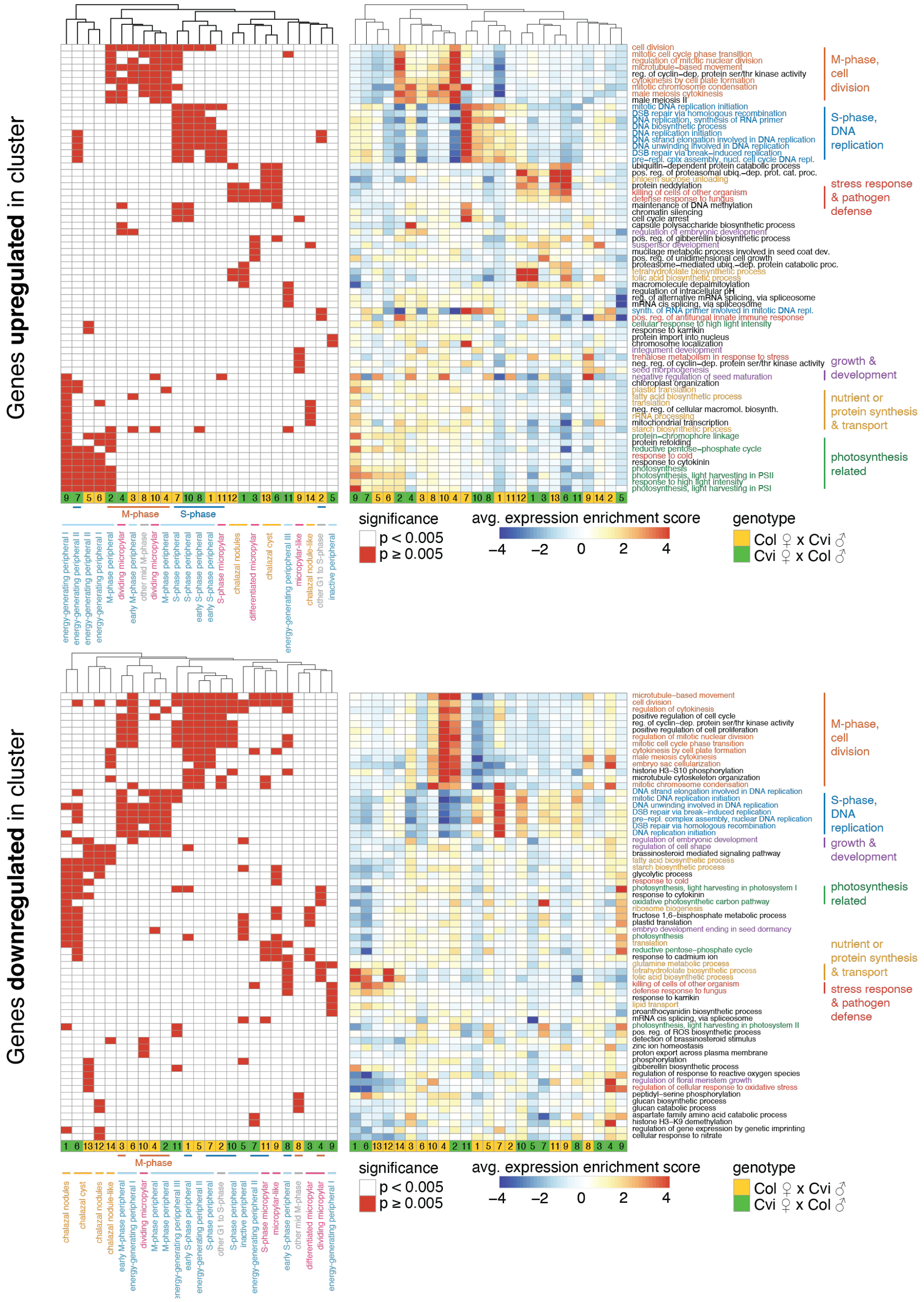


**Fig. S10. Heatmaps of the 5 most significantly enriched GO terms among genes upregulated (top) and downregulated (bottom) in each endosperm cluster.** Significant terms, $p < 0.005$, are flagged in left heatmap, while average expression ‘enrichment score’ across all genes associated with GO term is shown at right. Average includes any genes associated with the GO-term that are not significantly up/downregulated in the indicated cluster; so average may not reflect expectations. Order of rows/columns same for left and right heatmaps.

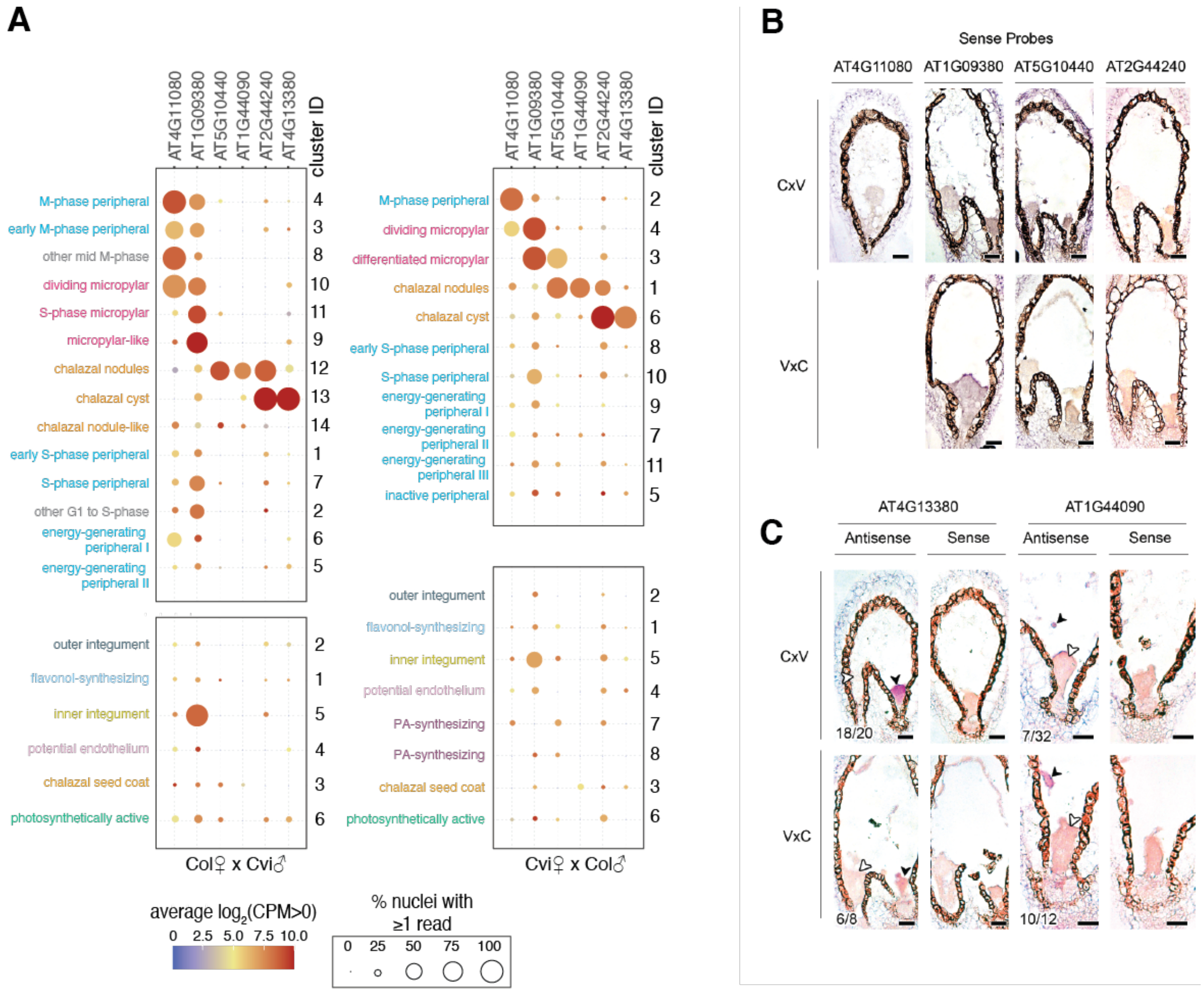


**Fig. S11.** ***In situ*** **hybridization analysis for additional cluster-specific transcripts and control sense probes.** (A) Expression data from snRNA-seq for all six marker genes used for RNA *in situ* hybridization experiments, across endosperm and seed coat clusters. (B) Sense probe images for probes used in Fig. 2. (C) *In situ* hybridization results for two additional transcripts not shown in Fig. 2: AT4G13380 is predominantly expressed in the chalazal cyst, while AT1G44090 is predominantly expressed in the chalazal nodules. Black arrowheads indicate sites of transcript accumulation; white arrowheads indicate sites without transcripts. Number of seeds with the pictured expression pattern, as well as total number of seeds observed, indicated in bottom left of each image. Both sense and antisense probe images shown.

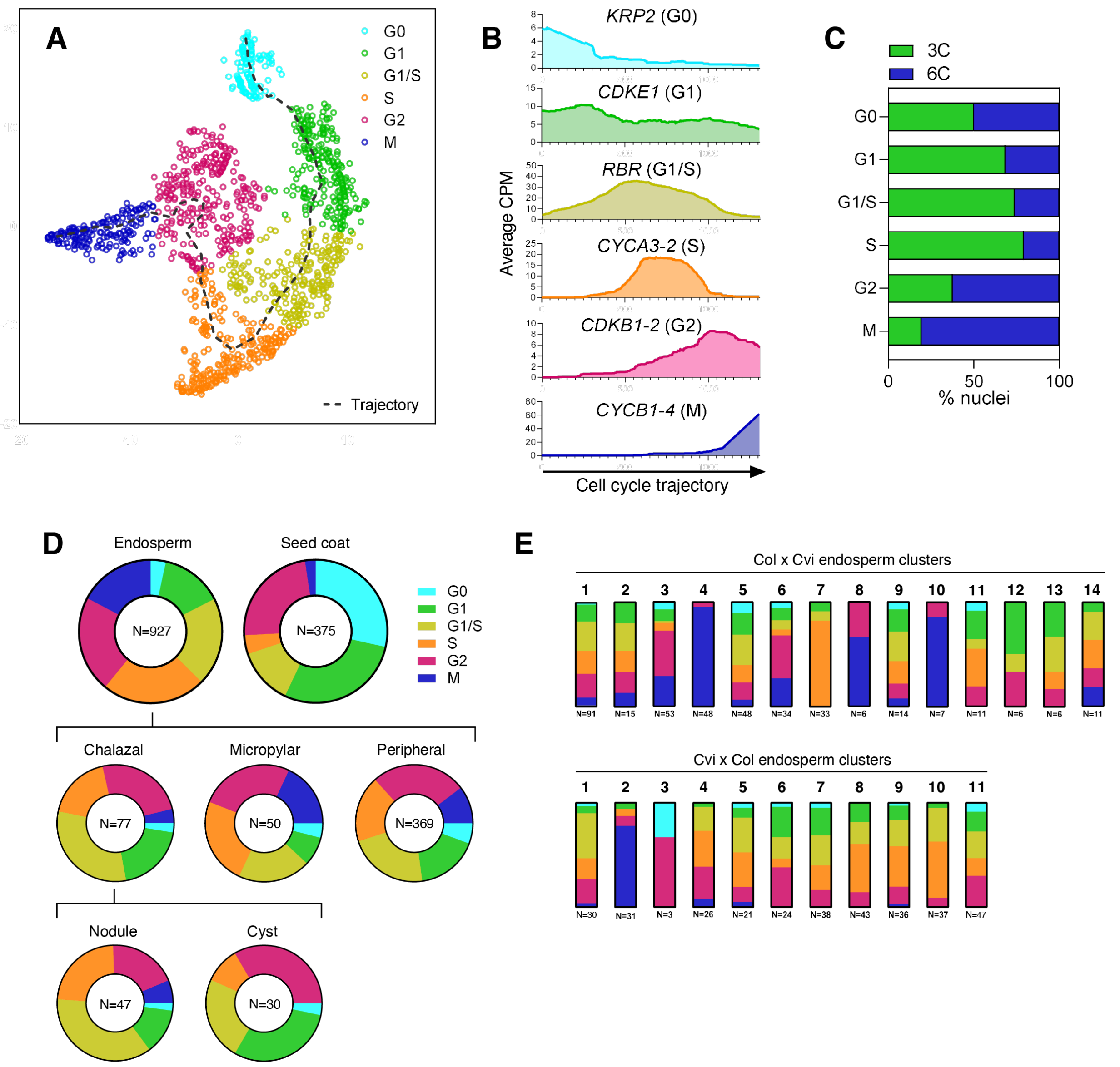


**Fig. S12. Cell cycle is a source of variability among endosperm clusters.** (A) t-SNE projection and trajectory analysis of 1,309 nuclei in the dataset, based on expression of a manually curated list of 22 cell cycle-dependent marker genes. Dotted line represents cell cycle trajectory from G0 -> G1 -> S -> G2 -> M. (B) Average expression of six of the 22 marker genes used in analysis shown in (A), with nuclei ordered according to their linear projection onto the cell cycle trajectory, starting from G0 (left) to M (right). Moving averages were calculated using a sliding window of 200 data points. (C) Percent of nuclei in each phase of the cell cycle that were sorted from the 3C or 6C FANS peak (D) Distribution of nuclei among cell cycle phases in seed coat and endosperm. Endosperm data are further divided into peripheral, micropylar, and chalazal; the chalazal region is also divided into the cyst and nodules. (E) Distribution of nuclei among cell cycle phases for each of the endosperm clusters.

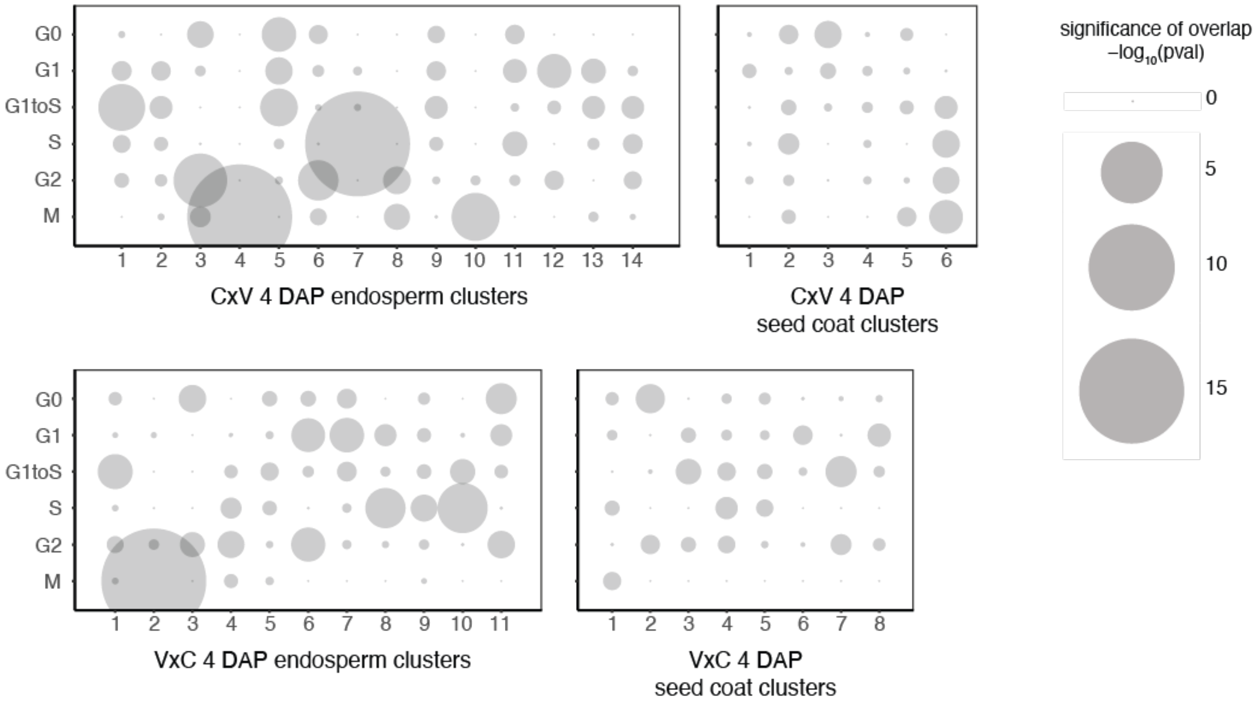


**Fig. S13. Significance of overlap between nuclei in each of the endosperm and seed coat clusters and nuclei in each stage of the cell cycle.** Cell cycle phase obtained from clustering analysis shown in Fig. S12. p-value obtained from hypergeometric test.

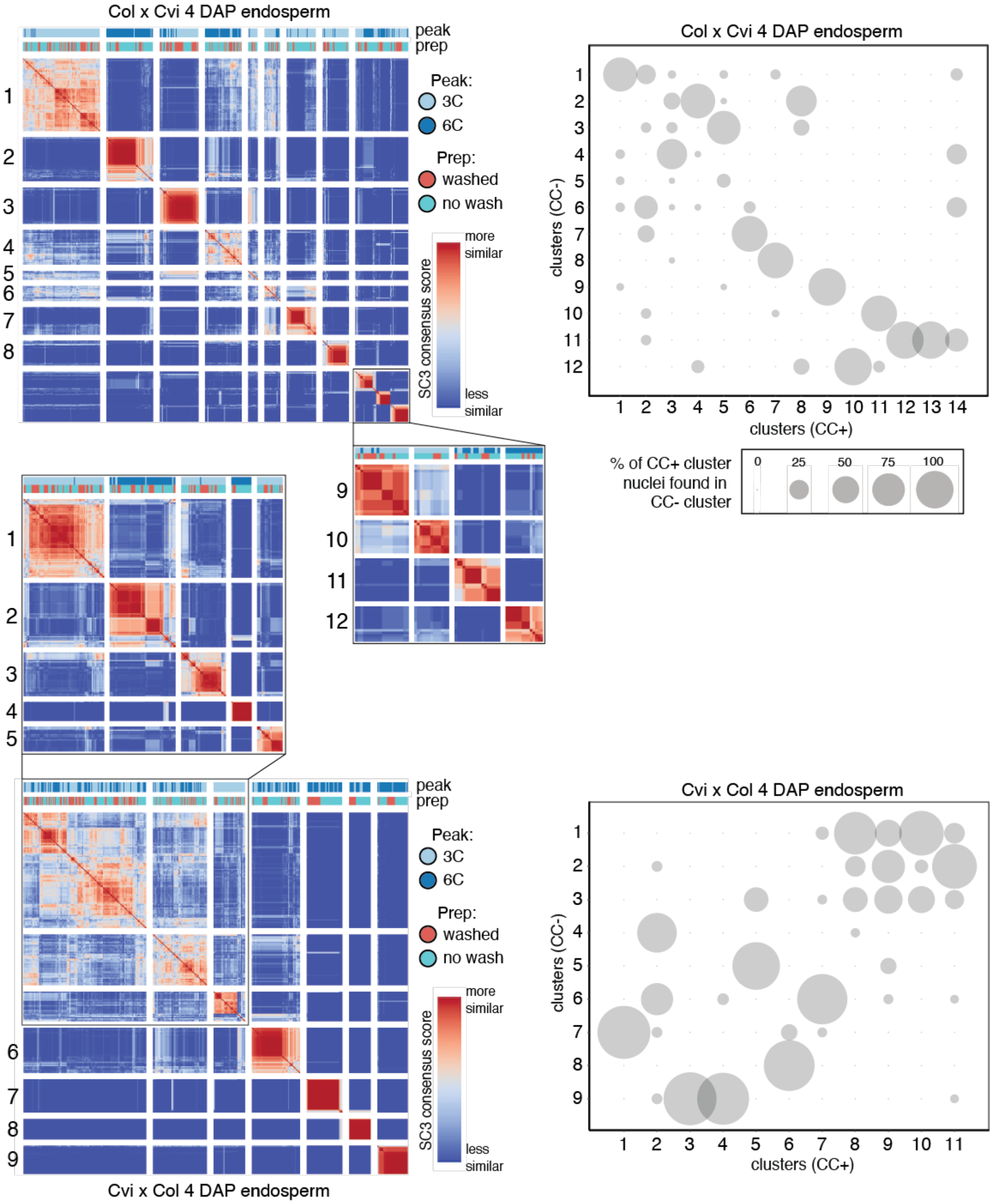


**Fig. S14. Omitting cell cycle correlated genes does not substantially alter clusters.** SC3 clustering was repeated while omitting the 1065 genes whose expression patterns correlated significantly with cell cycle trajectory. The nuclei in each of the new clusters obtained without cell-cycle correlated genes (CC-) were compared to those in the original clusters obtained from the full dataset (CC+), and the percent of CC+ nuclei that were present in the indicated CC- cluster are shown in plots at the right. Most nuclei from CC+ clusters fell largely (>75%) into a single CC- cluster, suggesting the clustering was largely unaffected by omitting the cell-cycle correlated genes. Clusters with strong expression of cell-cycle related genes (e.g. CxV E7; see Fig. 1D) were also not strongly affected.

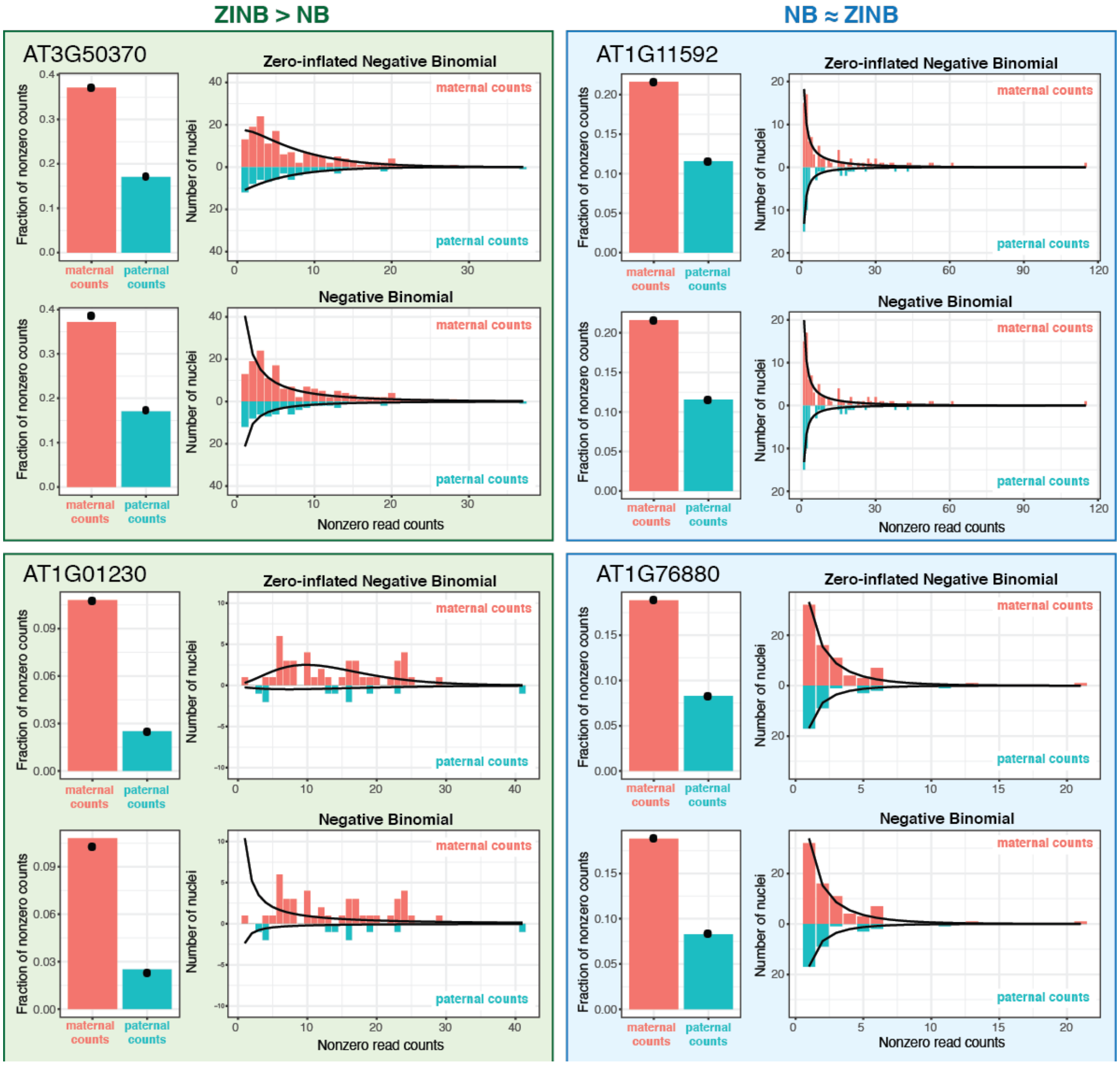


**Fig. S15. ZINB distribution accurately models allelic count data.** Examples of maternal and paternal count data from four genes fit to the zero-inflated negative binomial (ZINB) and negative binomial (NB) distributions. The fraction of nuclei with nonzero count values is given in bar chart to the left of each plot, with a histogram of the nonzero values on the right. Black dots on the bar chart and black lines in histogram indicate predicted values from ZINB (top) or NB (bottom) fit to the count data. On the left are genes where the ZINB distribution is better than the NB, on the right are genes where they are equivalent.

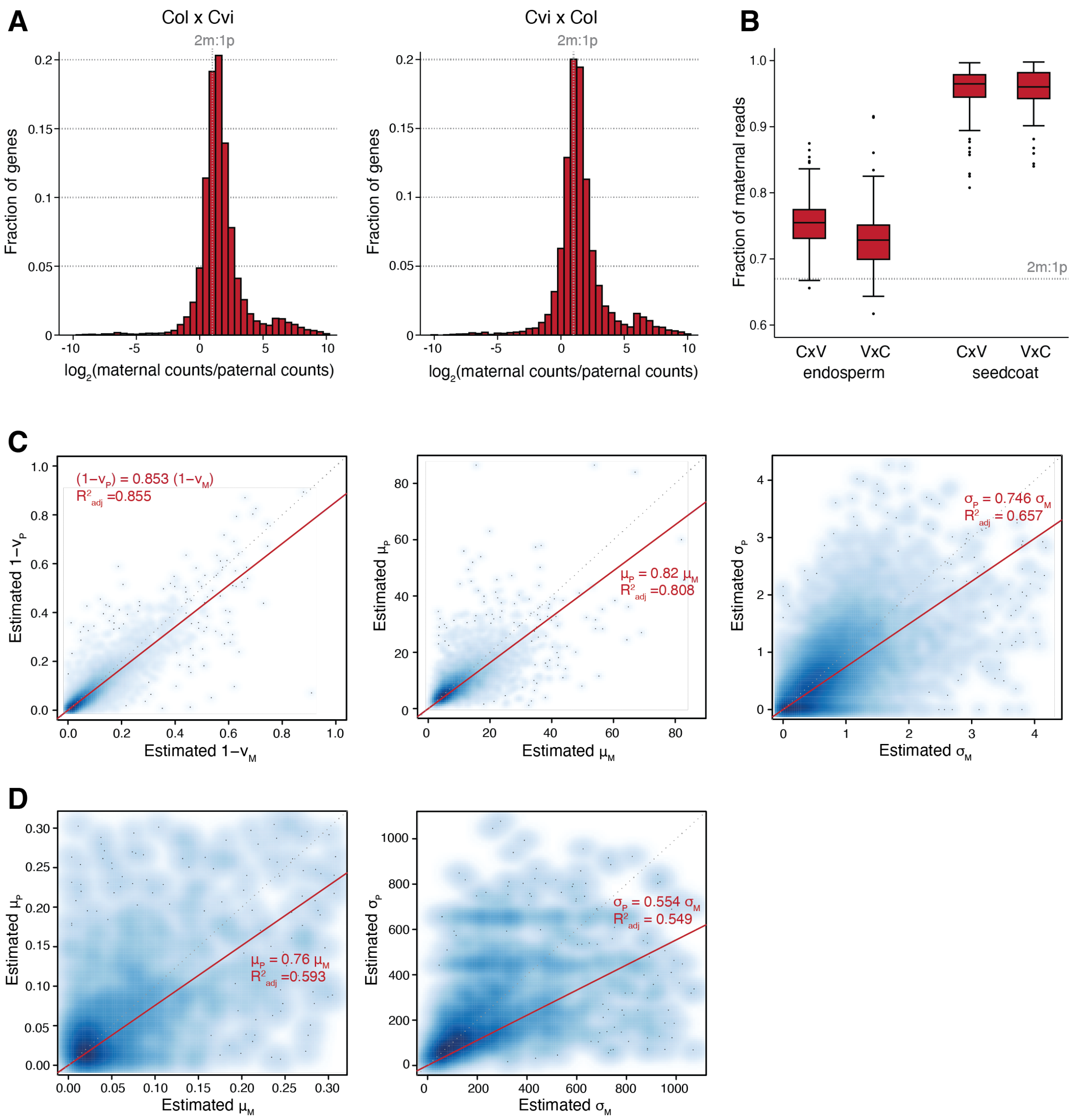


**Fig. S16. Exploration of maternal:paternal ratio and parameter estimates for fitted ZINB and NB models from the single-nuclei allelic count data.** (A) Histogram of the $\log_2$ ratio of maternal to paternal counts for all genes in the dataset, with the expected 2:1 ratio indicated as a grey vertical dotted line. (B) Fraction of maternal reads in CxV (Col x Cvi) and VxC (Cvi x Col) endosperm and seed coat. Maternal bias is more pronounced in CxV libraries than VxC, suggesting that the maternal bias is partially due to mapping bias in favor of the sequenced strain (Col). (C) Scatterplot of parameter estimates for the three ZINB parameters v, μ, and σ. Parameter estimates for maternal counts are plotted against estimates for paternal counts. Results shown from Col x Cvi. Each dot represents a single gene. (D) Scatterplot of parameter estimates for the two NB parameters μ and σ, for maternal and paternal count distributions.

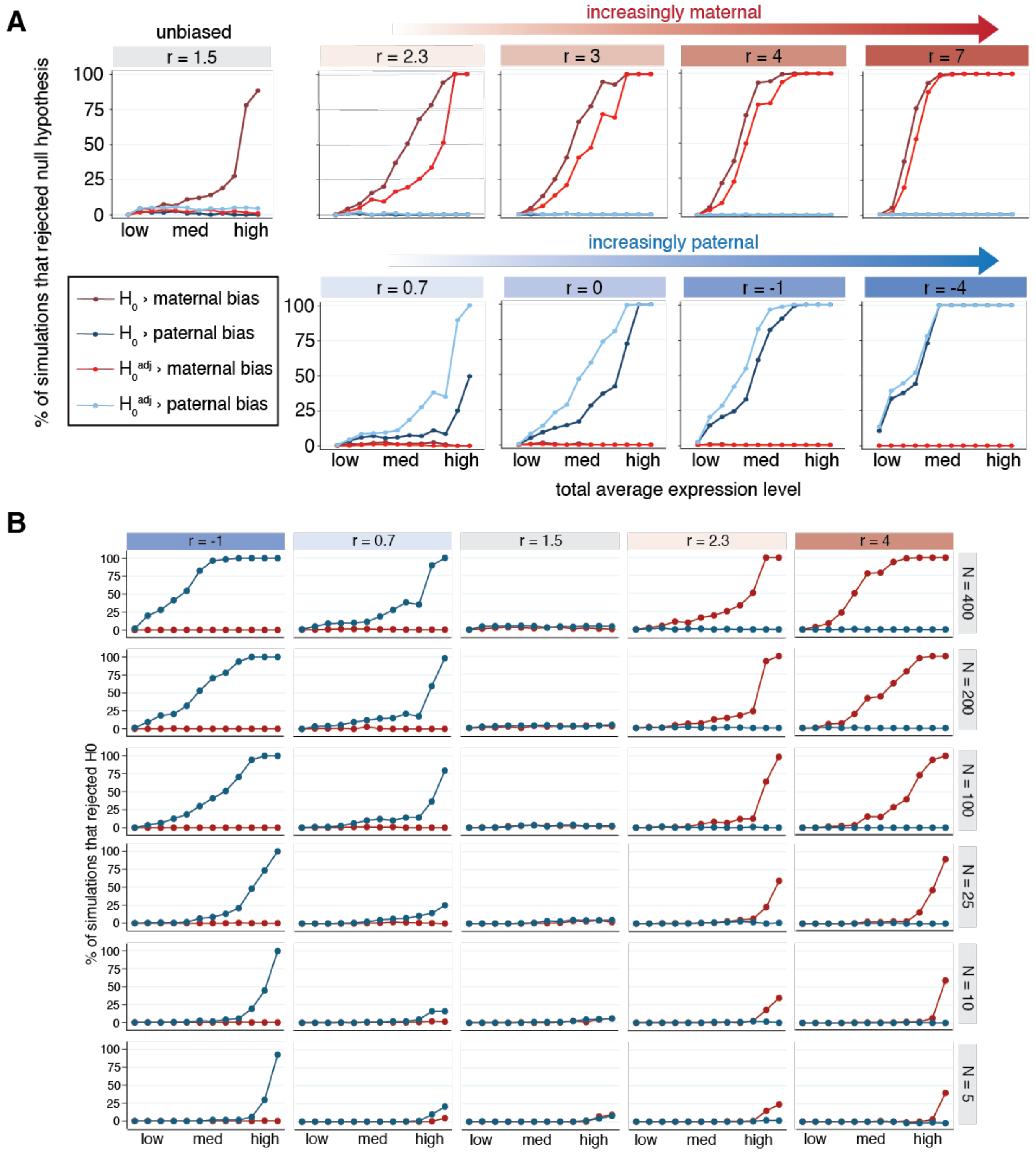


**Fig. S17. Statistical power and accuracy of imprinting model under various simulated conditions.** (A) Percent of simulations (out of 200) where the null hypothesis of no parental bias was rejected, for simulations with varied total expression and $log_2(m/p)$ ratio (r). Simulations mimicked degree of maternal skew in the Col x Cvi data, so 'unbiased' simulations had r = 1.5. Twelve values of total expression were tested: 0.01, 0.05, 0.1, 0.15, 0.25, 0.5, 0.75, 1.0, 1.5, 3.5, 15, and 50. The 1st, 25th, 50th, 75th and 99th percentiles for total expression in the Col x Cvi dataset are 0.033, 0.21, 0.58, 1.57 and 15.4, respectively. Blue lines indicate paternal bias, red indicate maternal bias. (B) Effect of number of observations (nuclei) in simulations on power to reject $H_0^{adj}$. Highly expressed and highly biased genes can be detected even with as few as 5 observations. Blue lines indicate tests for paternal bias, red indicate tests for maternal bias.

**A**

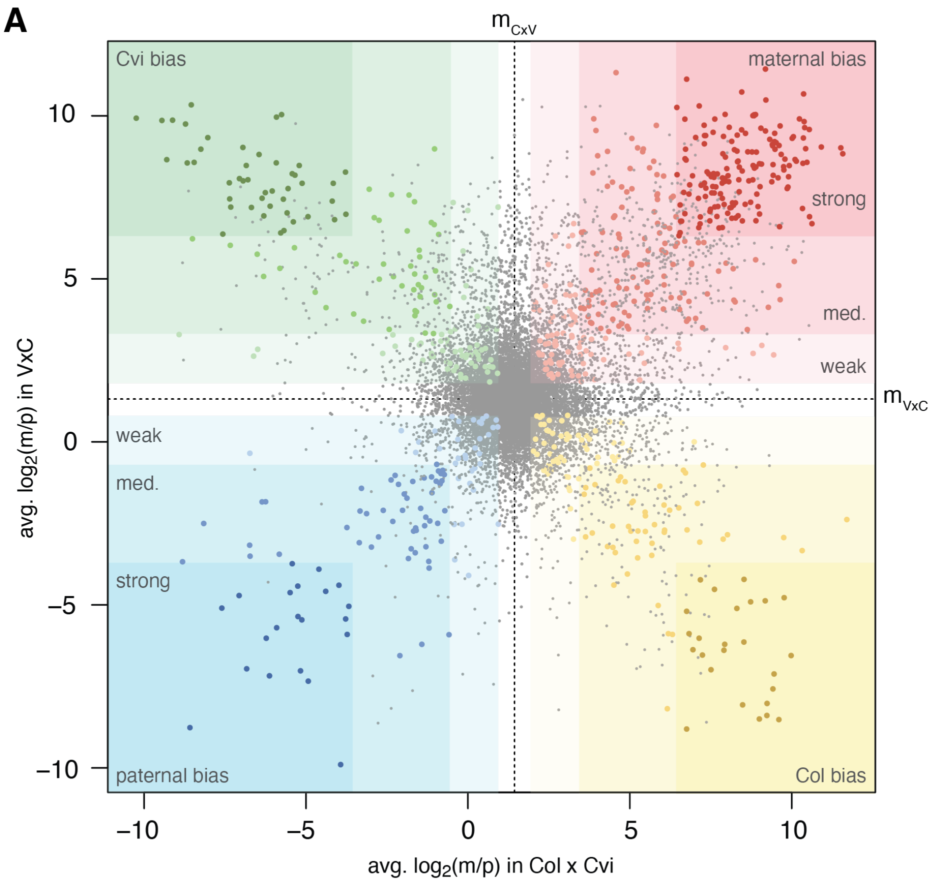


**B**

CxV status (rows) × VxC status (columns)

| CxV status \ VxC status | too few reads | failed to reject $H_0$* | weak Cvi bias | Cvi bias | strong Cvi bias | weak Col bias | Col bias | strong Col bias | total |
|---|---|---|---|---|---|---|---|---|---|
| too few reads | 18134 | 1328 | 1 | 12 | 27 | 13 | 38 | 70 | 19623 |
| failed to reject $H_0$ | 1284 | 11710 | 118 | 99 | 97 | 399 | 167 | 54 | 13928 |
| weak Col bias | 3 | 162 | 34 | 23 | 2 | 28 | 13 | 0 | 265 |
| Col bias | 17 | 113 | 17 | 55 | 39 | 16 | 31 | 6 | 294 |
| strong Col bias | 37 | 110 | 6 | 26 | 155 | 1 | 12 | 25 | 372 |
| weak Cvi bias | 5 | 368 | 28 | 11 | 3 | 20 | 9 | 1 | 445 |
| Cvi bias | 17 | 140 | 19 | 26 | 11 | 5 | 41 | 4 | 263 |
| strong Cvi bias | 29 | 59 | 3 | 10 | 45 | 1 | 8 | 21 | 176 |
| total | 19526 | 13990 | 226 | 262 | 379 | 483 | 319 | 181 | 35366 |

| | weak | med. | strong |
|---|---|---|---|
| too few reads | N/A | 18134 | N/A |
| failed to reject $H_0$ | N/A | 12757 | N/A |
| Col bias | 58 | 49 | 25 |
| Cvi bias | 64 | 47 | 45 |
| Col bias, CxV only | N/A | N/A | 223 |
| Col bias, VxC only | N/A | N/A | 221 |
| Cvi bias, CxV only | N/A | N/A | 199 |
| Cvi bias, VxC only | N/A | N/A | 196 |
| MEG | 82 | 120 | 155 |
| PEG | 36 | 53 | 21 |

**Fig. S18. Genes displaying allelic bias in reciprocal crosses of Col and Cvi.** (A) Scatterplot comparing weighted average $\log_2$(m/p) across all nuclei with allelic reads, in Col x Cvi compared to Cvi x Col. Quadrants of the plot corresponding to maternal, paternal, Col, and Cvi bias indicated. Genes that were significantly biased highlighted in corresponding color, with larger point size. Average $\log_2$(m/p) across all genes in Col x Cvi ($m_{CxV}$) and Cvi x Col ($m_{VxC}$) indicated by dotted black lines. Quadrants are divided into three subregions: weak bias (m ± 0.5), medium bias (m ± 2) and strong bias (m ± 5). (B) Table corresponding to (A), indicating number of genes in each category. Final classifications based on both Col x Cvi and Cvi x Col results, along with combined gene counts for each category, are shown in table at right.

**A**

| | Gene Status | # genes | % mat. CxV | % mat. VxC | $log_2$(m/p) CxV | $log_2$(m/p) VxC |
|---|---|---|---|---|---|---|
| MEGs | strong bias | 155 | 99.62 | 99.61 | 8.45 | 8.36 |
| MEGs | medium bias | 120 | 97.05 | 96.98 | 5.55 | 5.75 |
| MEGs | weak bias | 82 | 89.43 | 88.53 | 3.46 | 3.19 |
| PEGs | strong bias | 21 | 3.17 | 2.60 | -5.39 | -5.79 |
| PEGs | medium bias | 53 | 21.89 | 18.58 | -2.39 | -2.43 |
| PEGs | weak bias | 36 | 47.79 | 43.61 | -0.25 | -0.49 |
| Col bias | strong bias | 25 | 99.59 | 1.76 | 8.29 | -6.40 |
| Col bias | medium bias | 49 | 97.35 | 16.40 | 5.80 | -2.70 |
| Col bias | weak bias | 58 | 88.79 | 46.16 | 3.18 | -0.25 |
| Cvi bias | strong bias | 45 | 1.72 | 99.54 | -6.55 | 8.11 |
| Cvi bias | medium bias | 47 | 18.97 | 96.83 | -2.64 | 5.38 |
| Cvi bias | weak bias | 64 | 43.76 | 87.82 | -0.60 | 3.10 |
| other | CxV Col bias | 223 | 97.40 | 82.48 | 6.23 | 3.29 |
| other | CxV Cvi bias | 199 | 21.73 | 78.58 | -2.55 | 2.61 |
| other | VxC Col bias | 221 | 77.35 | 19.93 | 2.44 | -2.58 |
| other | VxC Cvi bias | 196 | 79.38 | 97.17 | 2.61 | 6.03 |
| no bias | | 12,757 | 71.60 | 69.88 | 1.56 | 1.43 |

**B**

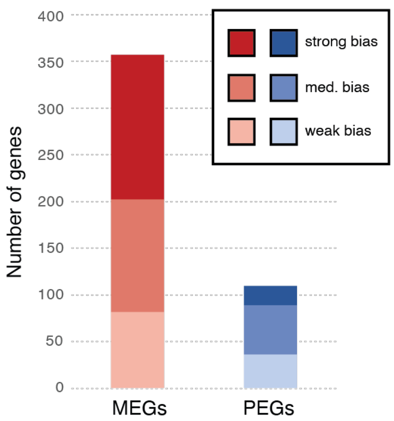


**Fig. S19. Summary of allelic bias under all conditions evaluated.** (A) Table of average percent maternal in CxV (Col x Cvi) and VxC (Cvi x Col) 4 DAP endosperm data, as well as average $log_2$(m/p) in each cross. (B) Number of MEGs and PEGs displaying strong, medium and weak bias (> 5, > 2, and > 0.5 s.d. away from median $log_2$(m/p) in dataset, respectively).

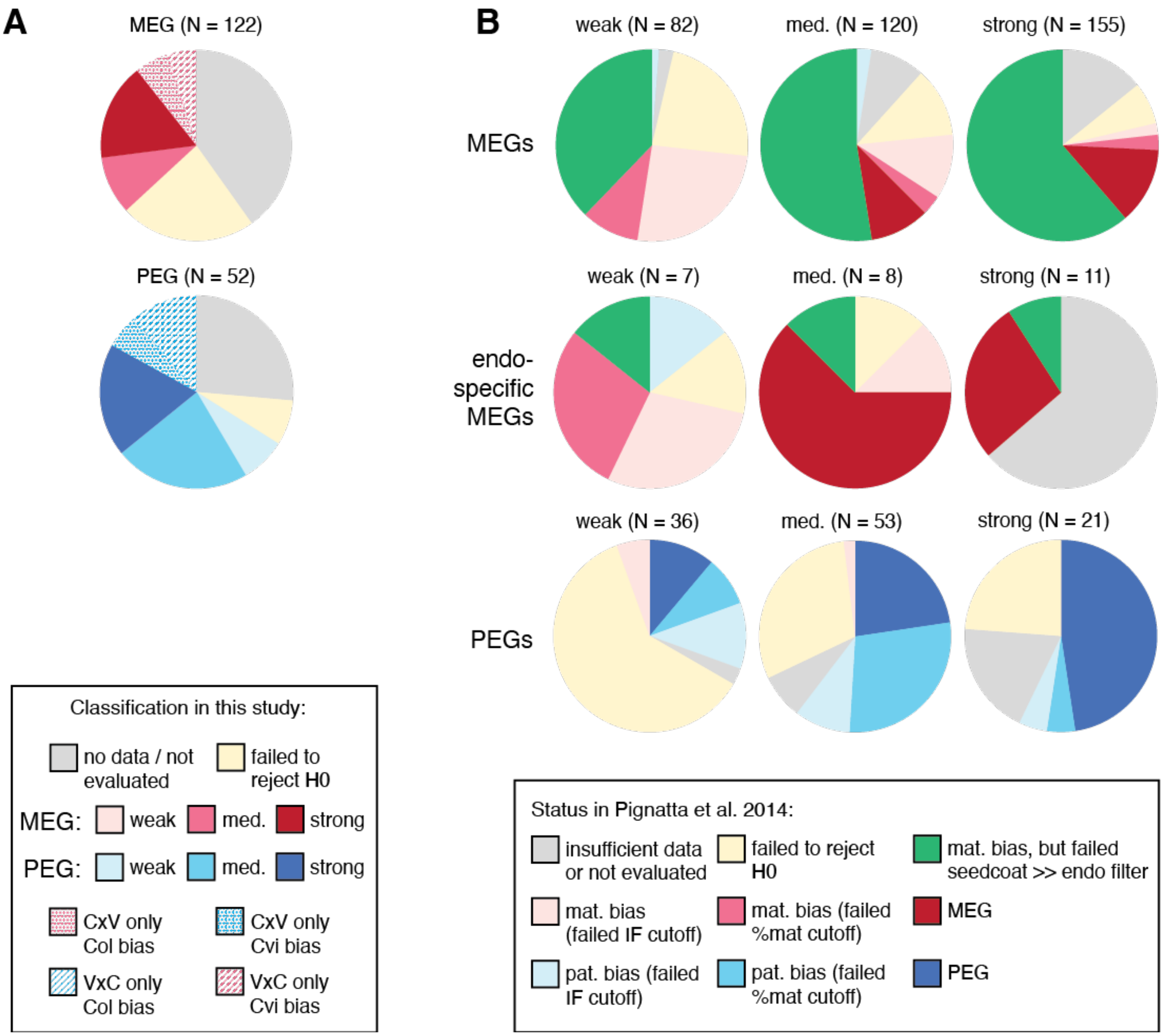


**Fig. S20. Comparison of imprinting status determined by snRNA-seq and conventional whole-endosperm mRNA-seq.** (A) Classification results from this study for the 122 MEGs and 52 PEGs identified between Col and Cvi 7 DAP endosperm in (*17*). Categories as shown in Fig. S18. For genes in the 'CxV only, Col bias' and similar categories, the null hypothesis could only be rejected for the indicated cross, but the other cross direction may also exhibit bias (but lack sufficient data to reject). (B) Status in (*17*) of all MEGs and PEGs identified in this study. (*17*) involved successive thresholds for assessing a gene's imprinting status: p-value < 0.01 -> IF (imprinting factor) > 2 -> % maternal either >85% (mat. bias) or <50% (pat. bias) -> imprinted (*34*). Genes that failed to meet all the imprinting criteria in (*17*) were therefore re-classified according to which cutoff they had failed. Additionally, maternally biased genes (p < 0.01) were subjected to an additional criterion that eliminated genes with much higher expression in seed coat vs. endosperm, to minimize potential biases that might be caused by seed coat contamination.

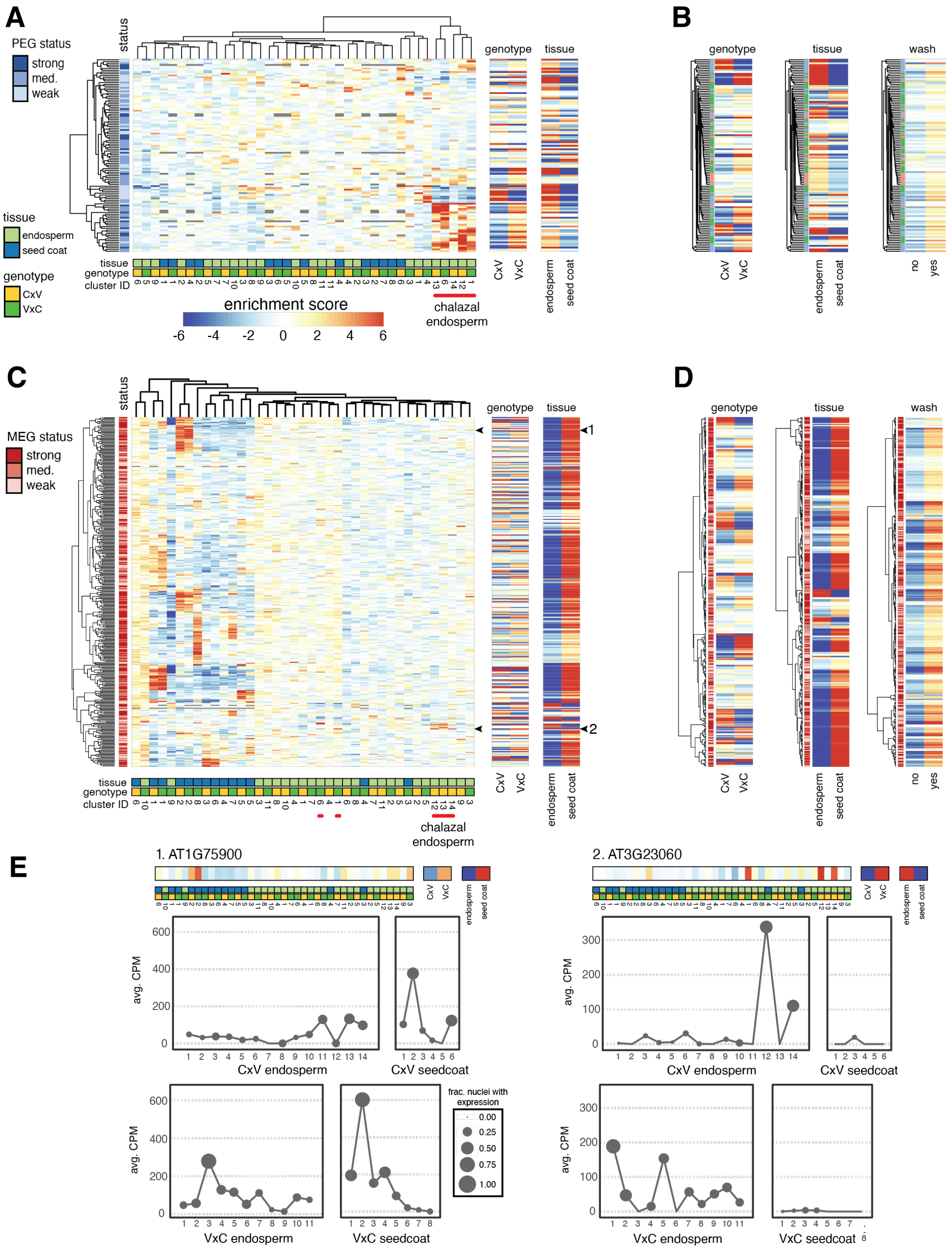


**Fig. S21. Summary of MEG and PEG expression patterns across endosperm and seed coat clusters.** (A,C) Hierarchical clustering of gene expression enrichment scores (ES) for all (A) PEGs and (C) MEGs, controlling for tissue and genotype. Separate heatmaps of ES calculated by genotype and by tissue on right. (B,D) Hierarchical clustering for (B) PEGs and (D) MEGs performed separately over ES from genotype, tissue and wash (yes = isolated nuclei were washed 2x prior to sorting, no = no extra wash steps). (E) Example expression profiles across nuclei clusters for two MEGs, AT1G75900 (strong MEG) and AT3G23060 (medium MEG). Locations of these genes in (C) indicated by black arrows.

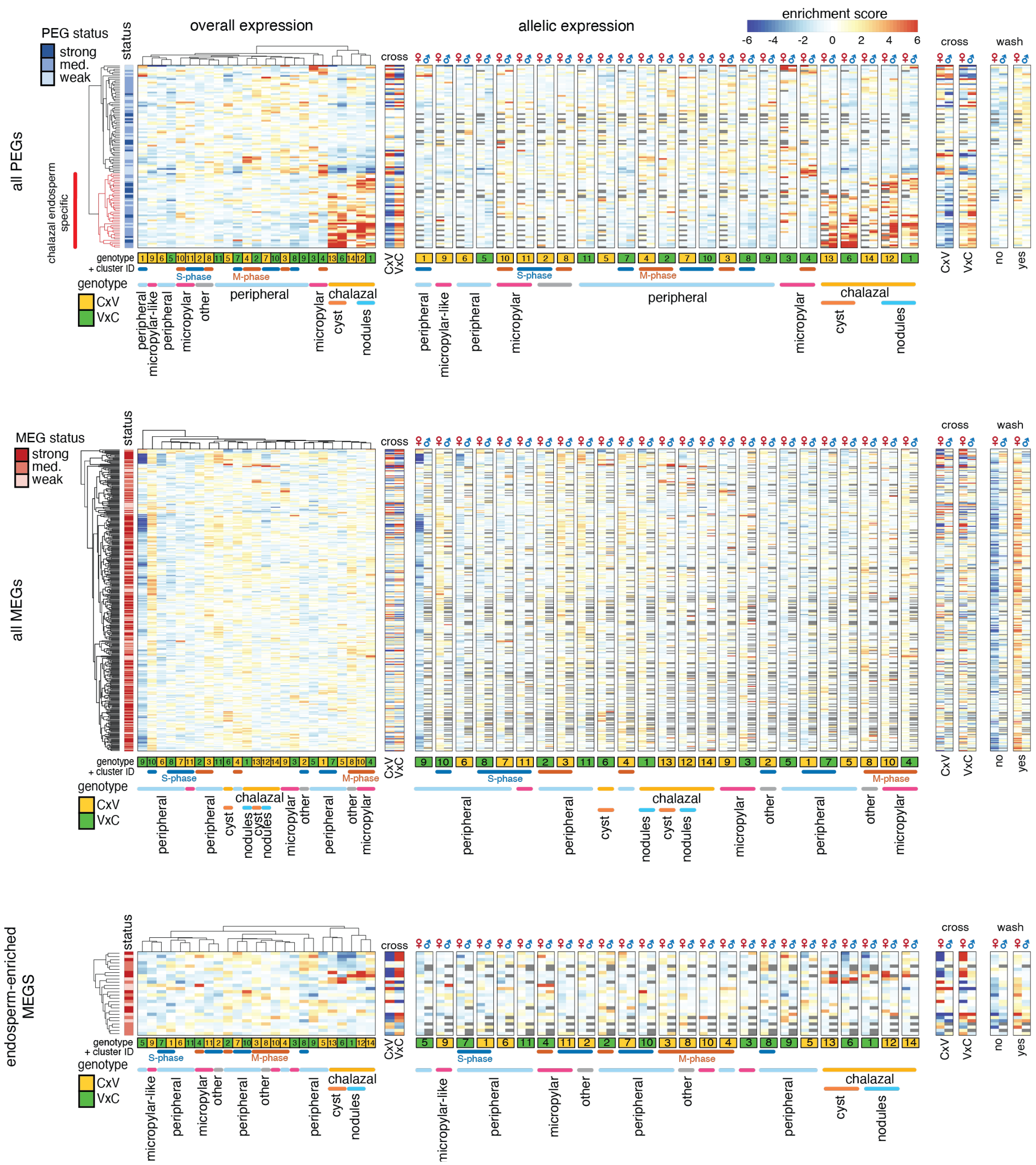


**Fig. S22. Total and allelic expression patterns across the endosperm clusters, for all imprinted genes.** Top panel shows all PEGs, middle all MEGs, and bottom panel shows a subset of MEGs whose expression is enriched in endosperm. Expression enrichment scores (ES) for total expression shown in heatmap on the left. On right, ES for maternal (♀) and paternal (♂) allelic expression enrichment scores are shown separately for each cluster, with row and column order otherwise identical to the leftmost heatmap. Scores were also calculated across factors cross (genotype of seed, CxV = Col x Cvi or VxC = Cvi x Col), and wash (nuclei were either washed 2x in sorting buffer prior to sorting, or not washed).

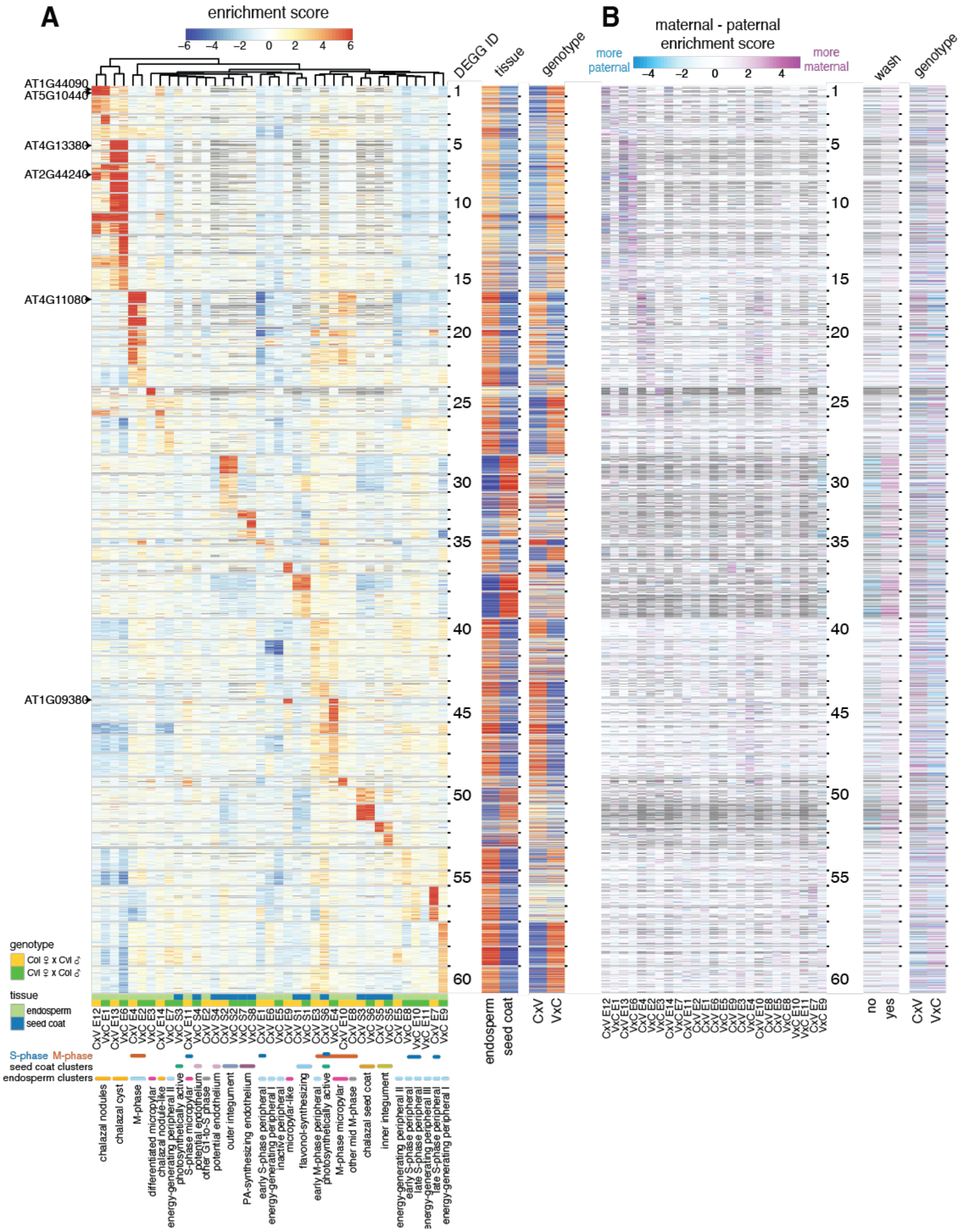


**Fig. S23. Total and allelic expression patterns of 4,500 genes significantly differentially expressed among the endosperm and seed coat clusters.** (A) Heatmap of gene expression enrichment scores (ES) (same as Fig. S7A). (B) Heatmap of difference between the maternal ($ES_{Mat}$) and paternal ($ES_{Pat}$) expression enrichment scores in the Col, Cvi 4 DAP endosperm dataset. Pink indicates relatively higher enrichment of maternal expression compared to paternal expression, while blue indicates the opposite. Far right: heatmaps comparing the $ES_{Mat}$ - $ES_{Pat}$ values as a function of sample prep (nuclei were either washed 2x in sorting buffer prior to sorting, or not washed) and genotype are also shown. Order of rows same in all plots.

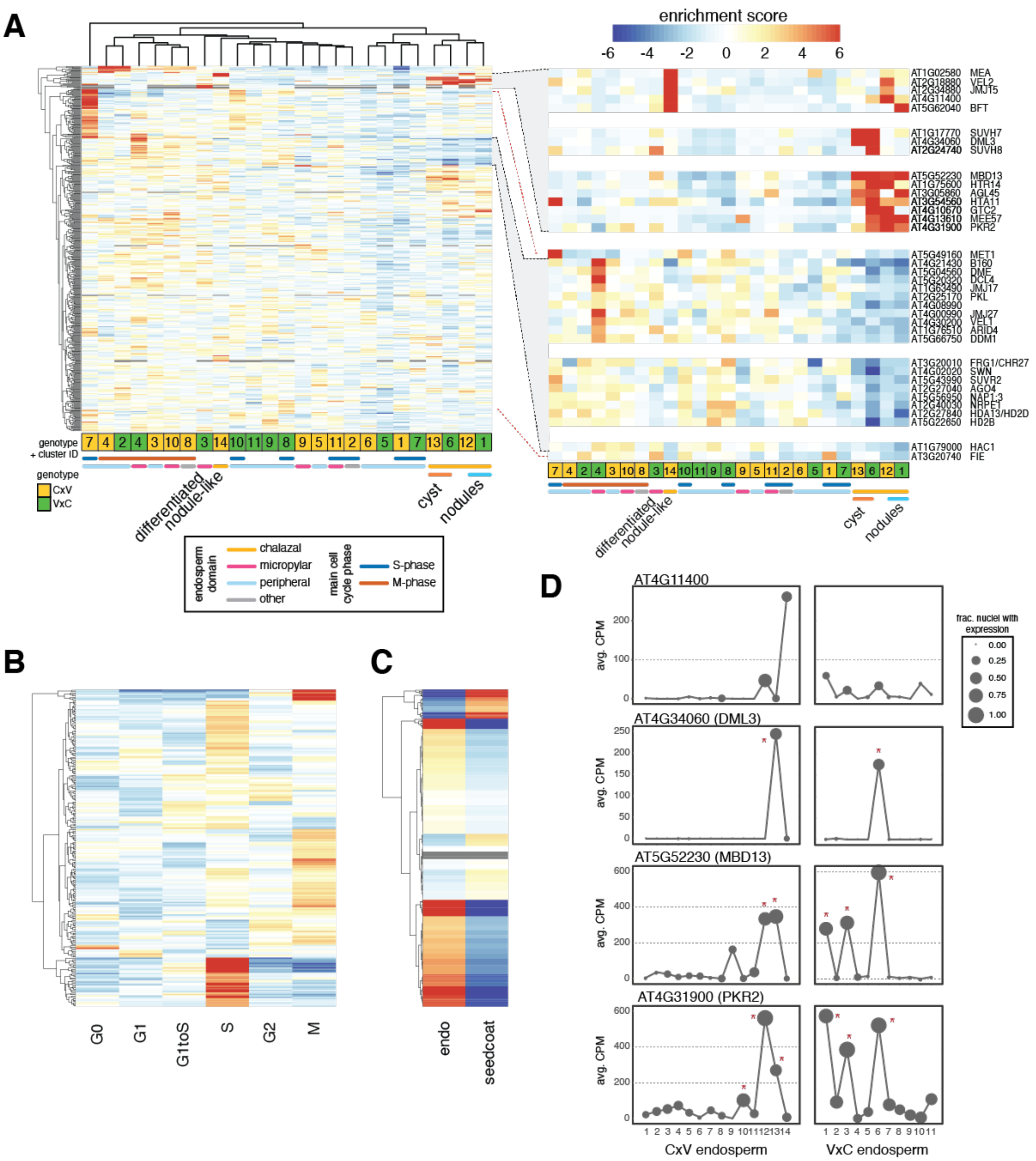


**Fig. S24. Expression of chromatin-related genes.** (A) Heatmap of expression enrichment scores (ES) across endosperm nuclei clusters, for 464 chromatin-related genes with variable expression across the clusters. Inset: subset of genes enriched in chalazal nodules, cyst, or both (top); subset of genes with depleted expression in chalazal endosperm, grouped by expression pattern (bottom). Not all genes in highlighted region in left plot shown. (B) Heatmap of expression ES for the full 4 DAP endosperm + seed coat dataset, over cell cycle phases. 227 chromatin-related genes with variation across cell cycle shown. Color bar same as (A). (C) Expression ES in endosperm vs. seed coat for 553 chromatin-related genes. Color bar same as (A). (D) Average expression profiles across the endosperm clusters for four genes shown in (A). Stars indicate clusters with significantly enriched expression based on a permutation test. CPM = counts per million.

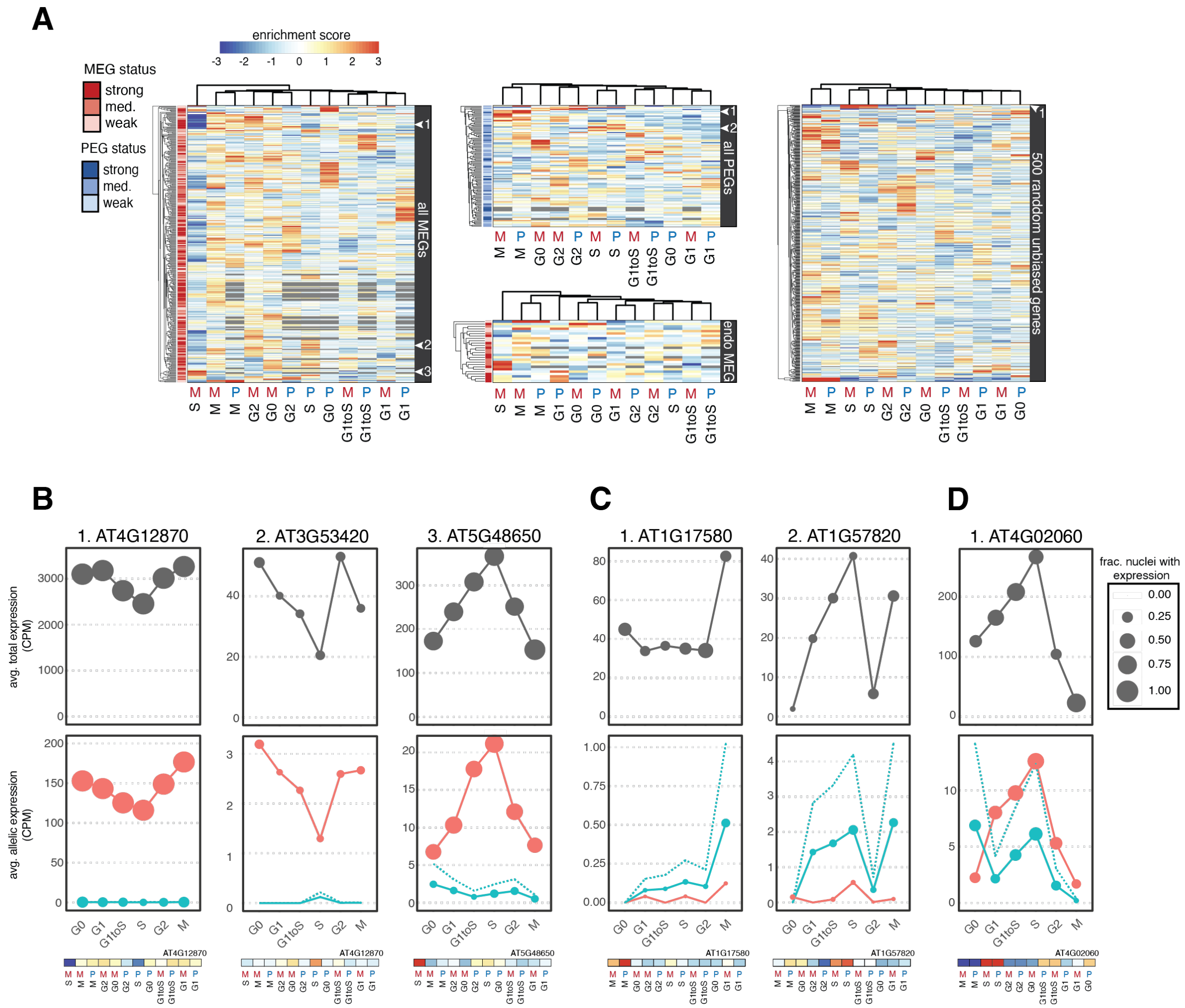


**Fig. S25. Effect of cell cycle on expression of imprinted genes.** (A) Enrichment scores for maternal (M) and paternal (P) allelic expression over different phases of the cell cycle, for all MEGs, all PEGs, MEGs specifically expressed in endosperm ('endo MEG'), and a random subset of 500 genes that show no overall allelic bias. (B-D) Example total (top, grey) and allelic (bottom, red = maternal, blue = paternal, dotted blue line = simulated 2x paternal) expression patterns across cell cycle phases for (B) three example MEGs, (C) two example PEGs and (D) an example non-imprinted gene. Genes in B-D are marked with white arrows in appropriate heatmap in (A). Values from heatmaps in (A) shown at bottom of each plot.
